# Competition among *Gardnerella* subgroups from the human vaginal microbiome

**DOI:** 10.1101/717892

**Authors:** Salahuddin Khan, Maarten J. Voordouw, Janet E. Hill

## Abstract

*Gardnerella* spp. are hallmarks of bacterial vaginosis, a clinically significant dysbiosis of the vaginal microbiome. *Gardnerella* has four subgroups (A, B, C and D) based on cpn60 sequences. Multiple subgroups are often detected in individual women, and interactions between these subgroups are expected to influence their population dynamics and associated signs and symptoms of bacterial vaginosis. In the present study, contact-independent and contact-dependent interactions between the four *Gardnerella* subgroups were investigated *in vitro*. The cell free supernatants of mono- and co-cultures had no effect on growth rates of the *Gardnerella* subgroups suggesting that there are no contact-independent interactions (and no contest competition). For contact-dependent interactions, mixed communities of 2, 3 or 4 subgroups were created and the initial (0 h) and final population sizes (48 h) were quantified using subgroup-specific PCR. Compared to the null hypothesis of additive interactions, most (69.3%) of the mixed communities exhibited competition (p < 0.0001). Competition reduced the growth rates of subgroups A, B and C. In contrast, the growth rate of subgroup D increased in the presence of the other subgroups (p < 0.0001). All subgroups were able to form biofilm alone and in mixed communities. Our study suggests that there is scramble competition among *Gardnerella* subgroups, which likely contributes to the observed distributions of *Gardnerella* spp. in vaginal microbiomes and the formation of the multispecies biofilms characteristic of bacterial vaginosis.

## Introduction

*Gardnerella vaginalis* is considered a hallmark of bacterial vaginosis, a dysbiosis of vaginal microbiome, although it is also commonly detected in women who do not meet the clinical criteria for vaginosis. *Gardnerella* comprises four sub-groups (A, B, C, and D), based on cpn60 barcode sequences and whole-genome sequences [1, 2]. These subgroups have also been classified as clades 1-4 [3], with subgroups A, B, C, and D corresponding to clades 4, 2, 1 and 3, respectively. More recently, Vaneechoutte et al. amended the description of *Gardnerella* and defined three new species within the genus: *G. leopoldii, G. swidsinskii*, and *G. piotti* [4]. These species correspond to three of the previously defined subgroups: *G. vaginalis* (subgroup C/clade 1), *G. leopoldii* and *G. swidsinskii* (subgroup A/clade 4), and *G. piotii* (subgroup B/clade 2), while subgroup D/clade 3 encompasses several unnamed “genome species”.

Phenotypic differences between the subgroups have been identified that could influence the role of *Gardnerella* spp. in the vaginal microbiome and their contributions to establishment and maintenance of vaginal dysbiosis [2, 4, 5]. Furthermore, it has been shown that clades 4, 1 and 3 (Subgroups A, C, and D) are more often associated with bacterial vaginosis as defined by a high Nugent score or Amsel’s criteria [6, 7]. Subgroup B or Clade 2 has been reported to be more abundant in women with an intermediate Nugent score [6–8]. Taken together, these observations highlight the potential clinical significance of the composition of the *Gardnerella* community within the vaginal microbiome.

*Gardnerella* clades or subgroups can be reliably distinguished in vaginal microbiome profiles using cpn60 barcode sequences [1, 9], whereas 16S rRNA gene sequencing does not provide sufficient resolution [4]. Profiling of vaginal microbiomes using cpn60 barcode sequencing, and application of clade-specific PCR has shown that the vagina is often colonized by multiple subgroups simultaneously [6, 7]. The proportional abundances of these subgroups, however, are not equal, and one subgroup usually dominates. The combinations and relative abundances of cpn60-defined subgroups of *Gardnerella* have been used to define previously undescribed population structures called community state types (CST) in the human vaginal microbiome [6]. Given the observed phenotypic diversity within *Gardnerella*, an understanding of the factors that determine *Gardnerella* population structure in the vaginal microbiome is critical.

Potential factors contributing to the abundance patterns of *Gardnerella* subgroups in the vaginal microbiome include differences among subgroups in terms of biofilm formation, adhesion, overall fitness, and resistance to anti-bacterial factors (either produced by other microbiota or delivered as a medical intervention). Interactions between the subgroups may influence the population dynamics of the *Gardnerella* subgroups [10–12]. When the vaginal microbiome is dominated by *Gardnerella*, interactions between subgroups would be more frequent than with other bacterial species because they are closely related and therefore more likely to occupy the same niche [13]. *Gardnerella* can form biofilm in isolation and can also be incorporated in multispecies biofilms in the vagina [14, 15]. Inter- and intraspecies interactions are ubiquitous within such multispecies biofilms [16–18], and such interactions may lead to competitive exclusion [19, 20]. Thus, it is possible that competition between *Gardnerella* subgroups within the biofilm shape the microbial population structure in the vaginal microbiome.

Competition between subgroups could take the form of a contest where two subgroups interact directly in either a contact-dependent or contact-independent manner involving the secretion of effectors that reduce the fitness of competitors [21]. Direct interactions can either inhibit the growth of one or more competitor(s) [22, 23], or trigger an enhanced biofilm response [19, 24]. In either case, competition could result in the exclusion of one or more competitor(s). Alternatively, competition between closely related taxa may take the form of a scramble [11], where they do not interact directly, but one competitor outgrows the others through its superior ability to use shared resources, such as nutrients. In a scramble mode of competition, all competitors have to share finite resources, which can cause reduction of the fitness of competing organisms. This type of competition is often referred to as non-interfering exploitative competition [25].

The objective of our present study was to seek evidence of contact-independent or contact-dependent interactions between *Gardnerella* subgroups that affect growth *in vitro*. Our results demonstrate that strains representing *Gardnerella* subgroups A, B, C, and D can coexist in biofilms but that mixing of subgroups does not enhance biofilm formation. Our findings also suggest the presence of a non-interfering, exploitative competition in mixed subgroups communities of *Gardnerella*.

## Methods

### *Gardnerella* isolates

Isolates of *Gardnerella* spp. used in this study were drawn from a previously described culture collection kept at −80 °C [2] (Table S1). The subgroup affiliations of all isolates were determined by cpn60 barcode sequencing [26]. Selected isolates were revived from −80 °C on Columbia agar plates with 5% sheep blood and incubated under anaerobic conditions (BD GasPak™ EZ Anaerobe Gas Generating Pouch System, NJ, USA) at 37 °C for 48 h.

### Measurement of the effect of *Gardnerella* culture supernatant on growth

The purpose of this part of the study was to test whether the *Gardnerella* subgroups produce molecules that inhibit the growth of the other subgroups (i.e. contact-independent interactions). Specifically, we wanted to test the effect of cell-free supernatant from *Gardnerella* subgroups on the growth and biofilm formation of the other subgroups. To detect contact-independent interactions, we tested a total of 56 combinations using 14 isolates of subgroups A (n = 4), B (n = 4), C (n = 3) and D (n = 3) (Table 1). Isolates that were used to derive cell-free supernatant (CFS) were called producer strains, and strains on which the effect of prepared CFS was tested were called focal strains. Selected focal strains of all four subgroups were grown in medium containing 10% CFS from producers belonging to other subgroups and their own subgroup (“self-CFS”). Experiments were performed in two culture media: NYC III broth, which is recommended by the American Type Culture Collection (ATCC) for *Gardnerella* culture, and BHI + 1% glucose broth, to determine if culture media influenced growth of *Gardnerella* subgroups in the presence or absence of CFS.

**Table 1.** Numbers of combinations of CFS producers and focal subgroups tested to detect contact-independent interactions

| Focal subgroup | CFS producer subgroup |  |  |  |
| --- | --- | --- | --- | --- |
|  | A | B | C | D |
| A | 3 | 3 | 3 | 3 |
| B | 4 | 4 | 4 | 4 |
| C | 4 | 4 | 4 | 4 |
| D | 3 | 3 | 3 | 3 |

To produce the CFS, colonies from Columbia blood agar plates were harvested using sterile swabs, resuspended in 5 ml of NYC III broth and incubated anaerobically for 72 hours at 37°C to reach stationary phase. CFS was generated by centrifuging the broth culture at 3 000 × g for 30 min (26). The supernatant was filter-sterilized using 0.22 μm filters and was used on the same day. Filter-sterilized CFS was streaked on Columbia blood agar plates to confirm sterility.

To test the effect of CFS on the focal strains, colonies from blood agar plates were harvested using sterile swabs, resuspended in 0.85% saline and adjusted to McFarland turbidity standard 1. Fifteen microliters of each test strain suspension were added to 135 μl of NYC III or BHI + 1% (v/v) glucose and 10% (v/v) CFS in individual wells of a flat bottom 96-well plate (Corning Costar, NY, USA). The focal strains were also grown in control wells with media containing no CFS. Negative controls consisted of 15 μl of 0.85% saline added to 135 μl of culture media (NYC III or BHI + 1% glucose) and sterile culture media alone. To confirm the viability of the inocula, focal strain suspensions were streaked on to Columbia blood agar. Each combination of CFS and the focal strain was performed in three technical replicates.

### Quantification of total growth, planktonic growth and biofilm growth

Total growth, planktonic growth, and biofilm formation were determined for each combination at 48 h (Fig S1-FigS14). Total growth was calculated as the difference in optical density measured at 595 nm between the 48 h and the 0 h time points. The OD_595_ was measured using a VMax Kinetic ELISA Absorbance Microplate Reader (Molecular Devices, CA, USA). Planktonic growth was measured by transferring 150 μl of supernatant from each well to a fresh 96-well plate and determining OD_595_. To measure the biofilm formation, a crystal violet (CV) staining assay was performed [19, 27, 28]. Briefly, after removal of the supernatant, plates were washed twice with water, biofilms were stained with 1% CV for 10 min, plates were washed twice with water and air dried. To solubilize stained biofilm, 150 μl of 33% glacial acetic acid was added to each well, and the OD_595_ was measured.

### Co-culture assays to detect contact-dependent interactions

The purpose of this part of the study was to test whether there were interactions between *Gardnerella* subgroups when they were grown together in the same culture. Four independent experiments were conducted at separate points in time with two different sets of *Gardnerella* isolates. Experiments 1A and 1B were done in March and April 2018, respectively, and subgroups A, B, C, and D were represented by isolates NR020, N170, NR038, and NR003. Experiments 2A and 2B were done in April and May 2018, respectively, and subgroups A, B, C, and D were represented by isolates VN003, VN002, NR001, and WP012. For each of the four replicate experiments, we grew the four subgroups alone (n = 4; A, B, C, D), and in all possible combinations of two (n = 6; AB, AC, AD, BC, BD, CD), three (n = 4; ABC, ABD, ACD, BCD) and four subgroups (n = 1; ABCD) for a total of 15 different combinations. Each of the 15 combinations was replicated three times in the wells of a 96-well tissue culture plate (i.e. 4 experiments *15 combinations * 3 replicates per combination = 180 replicates). The members of each community were allowed to interact for a period of 48 hours and the abundance of the constituent subgroups was estimated at the start (0 h) and the end (48 h) of this period using subgroup-specific quantitative real-time PCR (qPCR). Prior to the interaction assay, each of the four subgroups was grown alone at 37 °C anaerobically in BHI with 0.25% maltose and 10% horse serum for a period of 12 h and then mixed in BHI+ 0.25% maltose. Immediately prior to combining the subgroups to create the mixed communities, a sub-sample was taken from each of the four cultures to determine the abundance of each subgroup at the time point of 0 h using the subgroup-specific qPCR. To create the mixed communities, equal volumes of each isolate containing ∼5 × 10^6^ genome equivalents per mL (i.e., verified by qPCR) were included in a total volume of 200 µl per well. For each of the four experiments, two plates were used: one for quantifying the number of cells in both the planktonic and biofilm fractions using qPCR and the other to quantify biofilm formation using the CV staining assay.

### Quantification of *Gardnerella* using subgroup-specific quantitative real-time PCR

Cells from planktonic and biofilm fractions were collected at 48 h and extraction of DNA was performed using a commercial kit (DNeasy PowerBiofilm, Qiagen, Mississauga, ON) following the manufacturer’s instructions with minor modifications. To collect the planktonic phase, 200 μl of culture supernatant was removed from the wells and transferred to bead tubes. To collect biofilm, 200 μl of lysis reagent MBL was pipetted directly into the wells of the 96-well plate and incubated for 30 minutes to solubilize the biofilm. The bottom of each well was scraped with a pipette tip and the suspension was pipetted up and down several times before transferring it to a bead tube. The biofilm solubilization step was repeated to maximize biofilm collection. To enhance lysis, 100 μl of chaotropic agent FB was added to the bead tubes. The bead tubes were incubated at 65°C for 5 minutes in a water bath, and then vortexed using a multitube vortexer at maximum speed for 15 min. Later steps were performed following the manufacturer’s instructions.

Subgroup-specific quantitative real-time PCR was performed using previously published primers and probes [8] (Table S2). Amplification was performed in 10 μl reactions containing 1× iQ Supermix (BioRad, Mississauga, ON), 800 nM of each primer, 100 nM of TaqMan probe, and 2 μl of template. The qPCRs were performed using a CFX Connect (BioRad, Mississauga, ON) instrument. The qPCR results were reported as target copy number per PCR reaction (2 µl of template DNA extract). Each sample was assayed in duplicate reactions with the appropriate standard curve comprised of plasmids containing probe targets (10^2^ to 10^9^ plasmid copies per reaction). Thermocycling conditions were: initial denaturation at 95°C for 3 min, 40 cycles of 95°C for 15 s, and annealing/extension at 60°C (subgroups A, B, and D) or 63.3°C (subgroup C) for 40 s. Each plate contained a no template control, DNA extraction controls, and non-target subgroup templates as negative controls. For each qPCR reaction, the genome copy number was calculated using the standard curve. The qPCR assay was repeated for samples with a difference in Cq value > 1 between duplicate wells.

### Statistical analysis of contact-independent interactions

The contact-independent interactions were analyzed using Kruskal-Wallis nonparametric one-way ANOVA with Dunn’s post hoc test (Prism 8, Graphpad Software).

### Statistical analysis of contact-dependent interactions

To characterize the interactions between the four subgroups, the 180 replicates of 15 unique communities in four independent experiments (1A, 1B, 2A, 2B) were analyzed collectively (4 experiments * 15 combinations per experiment * 3 replicates per combination = 180 replicates; each of the 15 unique combinations was replicated 12 times). Outcomes from these co-culture experiments were interpreted as shown in Fig. 1 [29]. First, the growth rates of all isolates grown as singletons were calculated using the following formula: r_i_ = ln (N_t, i_/N_0, i_)/T. Here, r_i_ is the observed growth rate of subgroup i when it is alone (has no competitors), N_0, i_ and N_t, i_ are the initial and final population sizes of subgroup i as estimated by qPCR (sum of biofilm and planktonic cells), and T is the time period of 48 hours (Table S2). Under the null hypothesis of no competition, we used our estimates of r_i_ from the singletons and our estimates of N_0, i_ in the mixed communities to predict the expected abundance of each subgroup 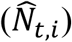 in the mixed communities after 48 hours of growth. The sum of predicted abundances (based on the singleton growth rates) for each subgroup in the community is the expected null abundance for the mixed communities (i.e. the expected community size in the absence of interaction). If the observed abundance of a mixed community was higher than the expected null abundance, the interaction was classified as a facilitation. If the observed abundance of a mixed community was lower than the predicted null abundance, it was classified as a competition (Fig. 1). The null hypothesis of no interaction predicts that due to random measurement error, 50% of the interactions should be positive (facilitation) and 50% of the interactions should be negative (competition). A proportion test was used to determine whether the observed prevalence of facilitation and competition were significantly different from the 50/50 expectation. This approach is a general test of the nature of interactions between *Gardnerella* subgroups and does not consider that each subgroup may be affected differently by competitors.

**Fig 1.**
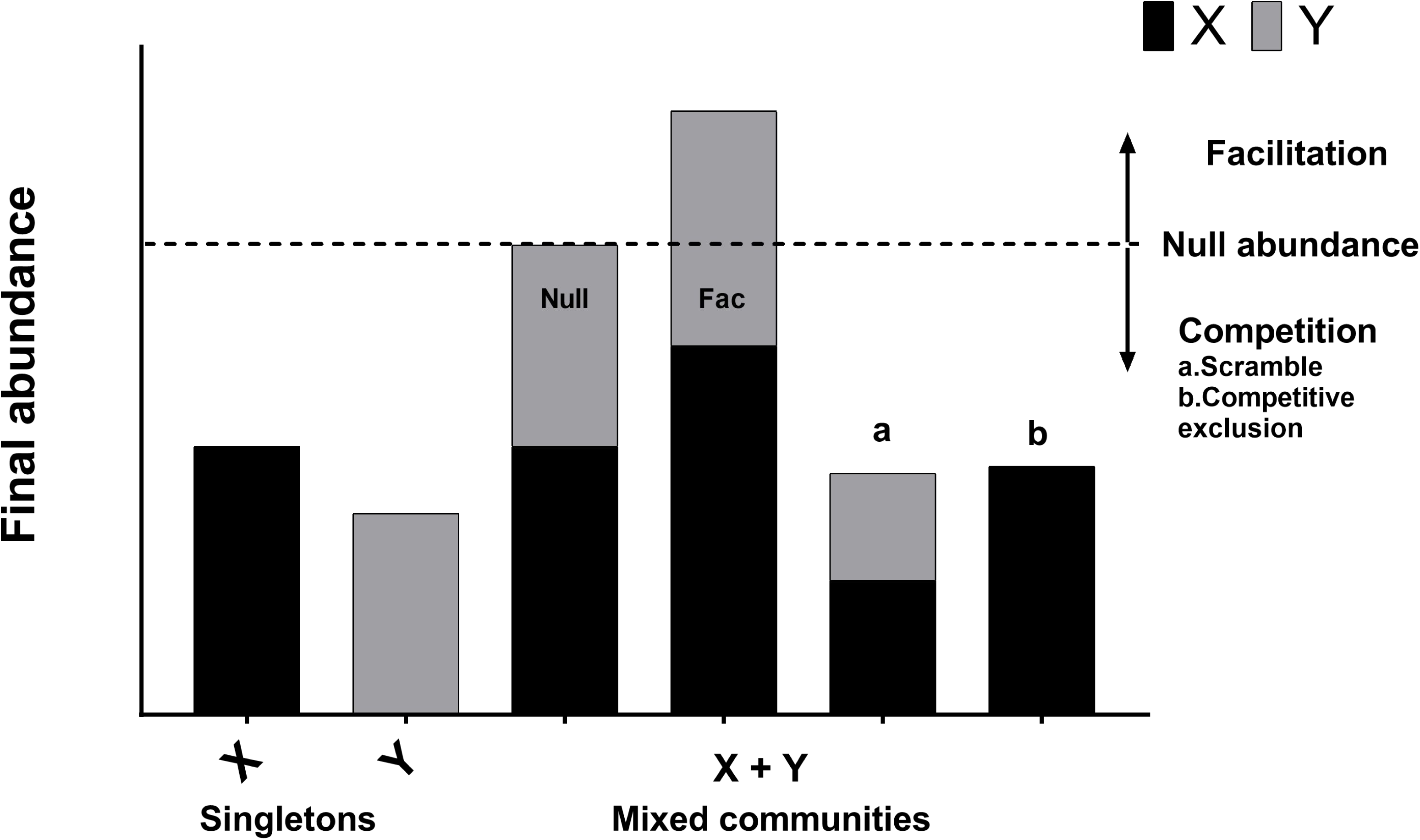
Regime of interpretation. Interactions between bacterial species or strains can be classified as neutral, competition, or facilitation. When bacterial species X and Y are grown in isolation, their abundances correspond to bars X and Y. Under the null hypothesis of no interaction, the expected abundance of the mixed community is equal to the sum of the abundances of two organisms grown separately (Null abundance, indicated by broken line). A neutral interaction occurs when the observed abundance of the mixed community is exactly equal to the null abundance (bar ‘Null’). In practice, neutral interactions are almost never observed due to measurement error. Facilitation occurs when the observed abundance of the mixed community is greater than the null abundance (bar ‘Fac’). Competition occurs when the observed abundance of the mixed community is less than the null abundance (bars ‘a’ and ‘b’). Competitive exclusion occurs when one of the species is completely eliminated in the mixed community (bar ‘b’).

To test whether the subgroups were affected differently by the number of competitors, the growth rates of the *Gardnerella* subgroups were analyzed using linear mixed effect models (LMMs). The residuals of the growth rates were considered as normally distributed. The fixed effects were subgroup (four levels: A, B, C, and D), the number of competitors (0, 1, 2, and 3), and their interaction. The random effects were the 3 replicates of each community nested in the 4 different experiments (i.e. total of 12 replicates for each community). This approach does not consider the identity of the competitors. We used R (v 1.1-21) to analyze the data; the LMM models were run using the lmer() function in the R package lme4.

To test whether the identity of the competitors mattered, the growth rate of each subgroup was analyzed separately using LMMs. The four subgroups had to be analyzed separately, because the identity of the competitors differs for each subgroup. The fixed effects were the identities of the competitors. For example, for the growth rate of subgroup A, the competitors included B, C, D, BC, BD, CD, and BCD. The random effects structure was the same as before.

## Results

### Effect of *Gardnerella* culture supernatant on growth and biofilm formation

The initial optical density (OD) of all the focal strains was ∼ 0.05 and they grew in both NYC III and BHI + 1% glucose, except for one subgroup C strain, NR001, which did not grow in BHI+ 1% glucose (Fig S11). The OD at 48 h of these strains varied from as low as 0.05 (after subtracting initial OD) to 0.80. There was no effect of CFS on overall growth or planktonic growth of focal strains, nor on biofilm formation (P > 0.05 for all comparisons) (Fig S1-S14). The type of medium, however, influenced mode of growth: NYC III had more planktonic growth, whereas BHI + 1% glucose had more biofilm growth. Increasing the concentration of CFS from 10% to 20% had no effect on the growth pattern of the *Gardnerella* subgroups (data not shown).

### Validation of qPCR assays

Prior to performing the co-culture experiments, the efficiency of each subgroup-specific qPCR assay and the limits of detection and quantification were determined, since these values had not been reported previously [8]. The amplification efficiency for subgroups A, B, C, and D were 99.9%, 107.4%, 110%, and 98.2%, respectively. The lowest concentration at which all subgroups were detected was 1 target copy per qPCR reaction. However, the lower limit of quantification (LOQ) was different for each subgroup. The LOQ for subgroups A, B, C and D were 1, 10, 100, and 1 copy per reaction, respectively.

### Characterization of contact-dependent interactions between *Gardnerella* isolates

Our null hypothesis approach of testing the type of interaction (facilitation versus competition) between subgroups of *Gardnerella* found that competition was 2.3 times more common than facilitation. Of the 132 mixed communities, 69.7% (92/132) had negative interactions (competition), and 30.3% (40/192) had positive interactions (facilitation). A proportion test found that these percentages were significantly different (p <0.0001) from the 50/50 null expectation. Competition was more frequently observed in communities with more subgroups. The prevalence of competition was 58.3% (42/72), 79.2% (38/48) and 100.0% (12/12) in communities with two, three, or four subgroups, respectively.

### Competition between subgroups in biofilms versus the supernatant

Since biofilms are a common site of interactions between species [16, 17, 19, 30], we investigated whether competition was more frequent in the biofilm fraction than the planktonic fraction. Out of 132 mixed communities, 68.9% of the biofilm fractions (91/132) and 65.9% (87/132) of the planktonic fractions exhibited competition. This result indicates that competition occurs in both biofilm and planktonic populations of *Gardnerella* spp. In addition, these observations demonstrated that mixed subgroup biofilms can occur, with no subgroup excluded.

### Effect of mixed communities on the growth rate of individual subgroups

When grown in isolation, the instantaneous growth rates (per hour) of the four subgroups ranked from lowest to highest are as follows: 0.098 for subgroup A, 0.119 for subgroup D, 0.120 for subgroup B, and 0.211 for subgroup C. The population doubling times (in hours) ranked from slowest to fastest are as follows: 7.1 for subgroup A, 5.8 for subgroup D, 5.8 for subgroup B, and 3.3 for subgroup C. The LMM found a significant interaction between subgroup and number of competitors on the growth rate (p <0.0001) indicating that the effect of the number of competitors on the growth rate differs between subgroups. Growth rates of subgroups A, B, and C decreased significantly (p <0.0001) in mixed communities (Fig. 2a, 2b, 2c). In contrast, the growth rate of subgroup D increased significantly (p <0.0001) in mixed communities (Fig. 2d). Thus, subgroups A, B, and C experienced competition in mixed communities, whereas subgroup D experienced facilitation. Regardless of the identity of the community, subgroup C always has a higher intrinsic growth rate than the other subgroups (Fig. 2).

**Fig 2.**
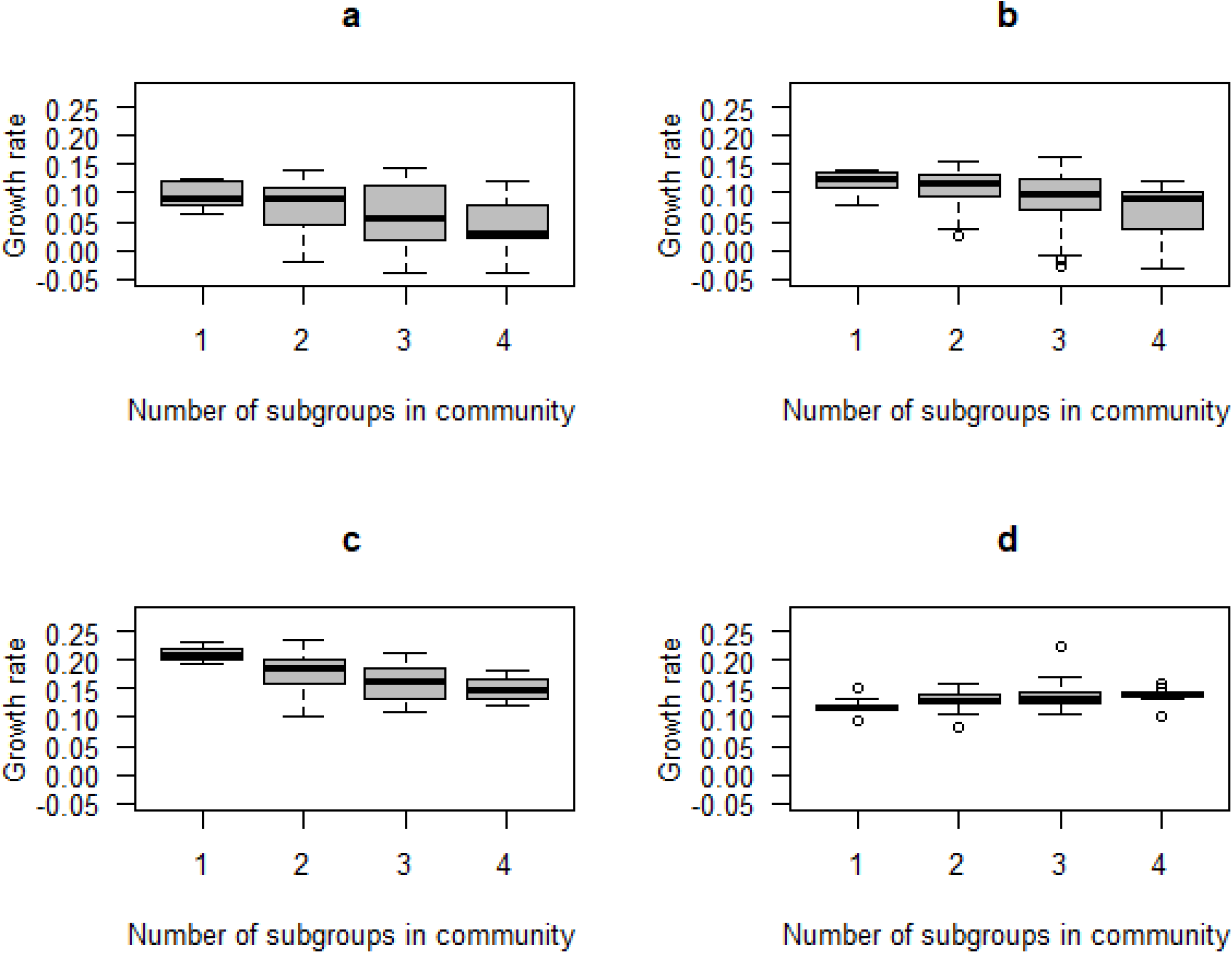
Effect of mixed communities on growth rates of *Gardnerella* subgroups. The growth rates of subgroups A, B, and C decreased with an increasing number of competitors indicating competition (panels a, b, and c). In contrast, the growth rate of subgroup D increased with an increasing number of competitors indicating facilitation (panel d). Each community was allowed to interact and grow over a period of 48 hours.

### Impact of competitor subgroups on focal subgroups

Next, we investigated whether subgroups differed in the magnitude of their negative (or positive) effect on the growth rate of other subgroups. Subgroup D had the most negative impact on the growth rates of the other subgroups. The presence of subgroup D in any community reduced the growth rate of the other members of the communities by 44.2% (Fig. 3d; p <0.0001). Subgroup A reduced the growth rates of the other members of the communities by 4.8%, whereas B and C increased the growth rates of the other communities by 7.2%, and 1.6%, but none of these effects were statistically significant (Fig. 3a, 3b, and 3c). In summary, subgroup D has a large and negative effect on the growth rate of all other subgroups, whereas the effects of the other subgroups are essentially neutral.

**Fig 3.**
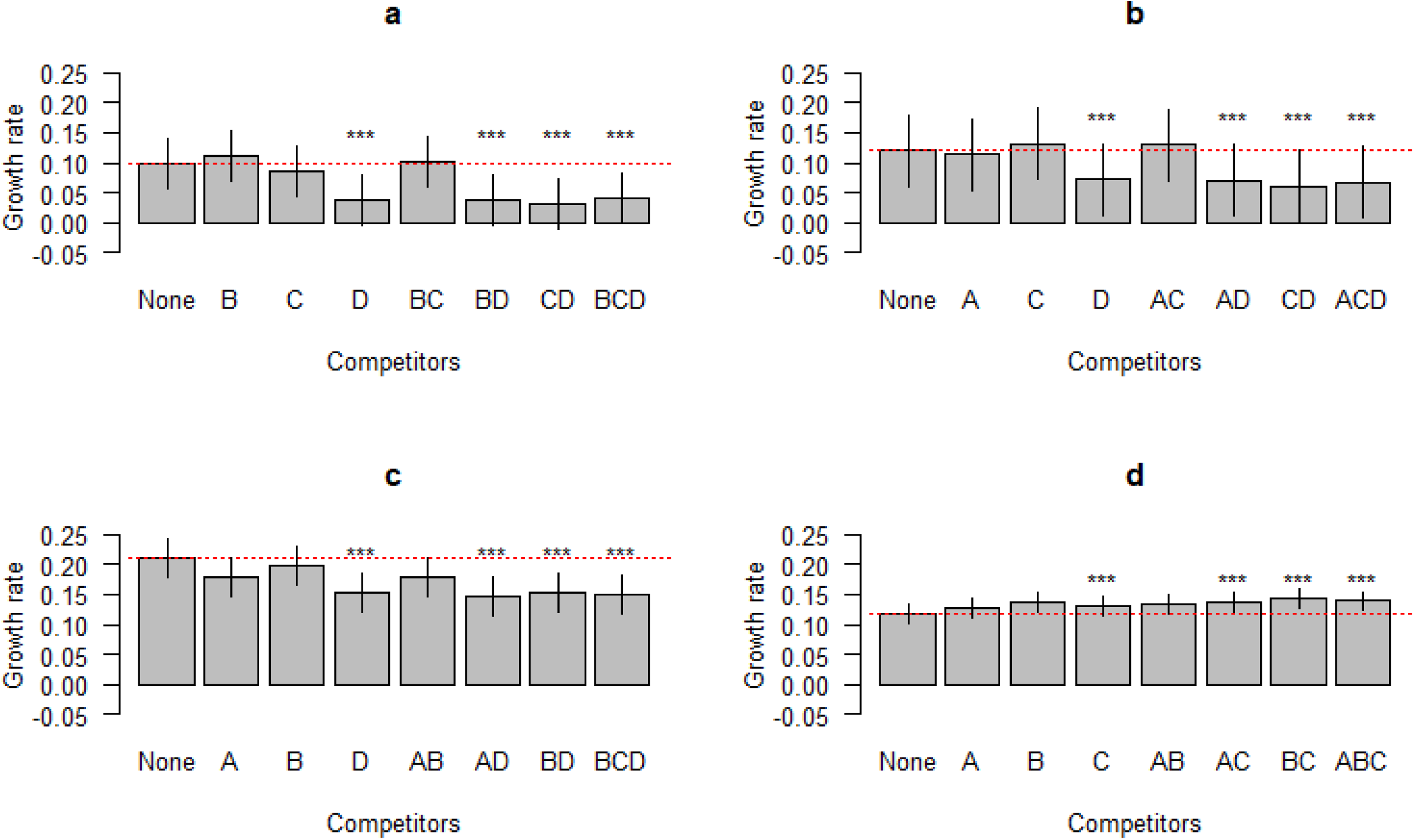
Impact of competitor subgroups on focal subgroup. The growth rates of subgroups A, B, and C are inhibited by the presence of subgroup D (panels a, b, and c). In contrast, the growth rate of subgroup D is enhanced by the presence of the other subgroups (panel d). The identities of the competitors are indicated on the X axis. The red dotted line is the expected null growth rate (growth as a singleton). Growth rates above and below the red dotted line indicate facilitation and competition, respectively. Significant effects on growth rates of the focal subgroups in the presence of competitors are denoted by asterisks (***). The vertical lines on each bar indicate the 95% confidence intervals for the means.

### Quantification of biofilm formation in monocultures and co-cultures

Mixing of different bacterial species often leads to increased biofilm formation. We therefore investigated whether mixing of *Gardnerella* subgroups would enhance biofilm formation. If the amount of biofilm formed by a mixture of subgroups was greater than the amount formed by the best individual biofilm former of that mixture, the interaction was considered synergistic. If the amount of biofilm formed by a mixture was less than the amount formed by the worst individual biofilm former of that mixture, the interaction was considered antagonistic [31]. Here biofilm formation by mixed subgroups was almost always less than the individual biofilm formation by the best biofilm-forming subgroup (Fig. 4 a, b). Only one co-culture of subgroups A and D was higher than the individual biofilm formation of both strains (Fig. 4b). A proportion test found that these proportions were not significantly (p >0.05) higher than 50/50 null expectation. The results of this experiment show that overall biofilm biomass is not enhanced by mixing of subgroups.

**Fig 4.**
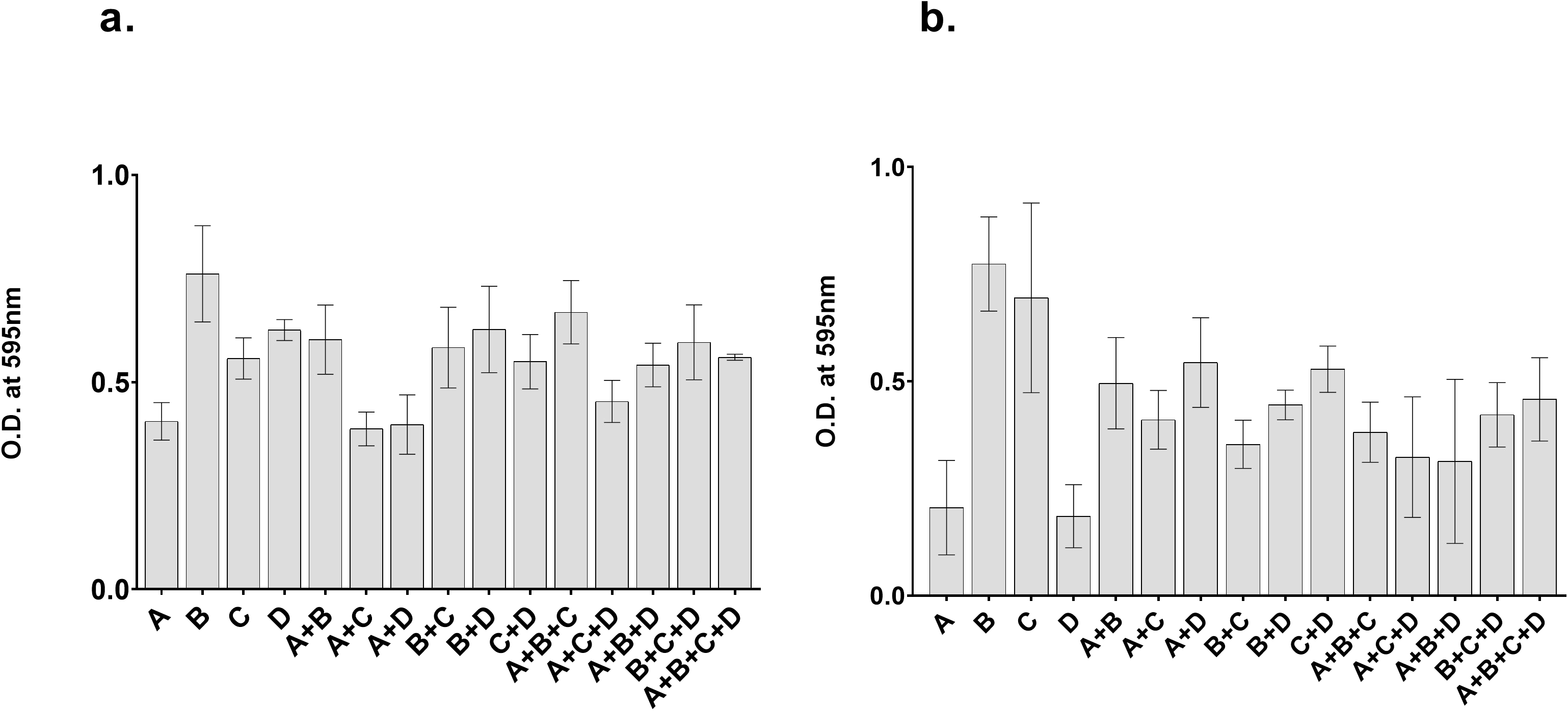
No enhancement of biofilm formation in mixed communities. Biofilm biomass was quantified by a crystal violet assay after 48 hours of growth. The error bars indicate standard deviations of three replicates.

### Effect of *Gardnerella* co-culture supernatant on individual subgroups

Our initial experiments showed that the CFS of singleton cultures had no impact on the growth of isolates from other subgroups, but negative interactions were frequently observed in co-cultures. To determine if effectors were secreted as a result of contact, we derived CFS from pairwise co-cultures and prepared media conditioned with 10% co-culture supernatant. We grew four representative isolates of all four subgroups in media with and without co-culture CFS and measured optical density to monitor growth and used a CV assay for quantification of biofilm formation (Fig S15). No significant differences in the amount or mode of growth were observed with exposure to co-culture CFS (p > 0.05).

## Discussion

A critical step in the development of bacterial vaginosis is when *Gardnerella* spp. displace *Lactobacillus* spp. and initiate multispecies biofilm formation [32]. The recent amendment of the genus and the “fine-tuning” of the taxonomy of *Gardnerella* [4] re-emphasizes the clinical importance of determining the particular contributions of different *Gardnerella* spp. to vaginosis, and the degree to which different subgroups may compete or cooperate in the vaginal microbiome. The detection of multiple subgroups (or species) of *Gardnerella* in the vaginal microbiomes of individual women is common and so interactions are expected to occur frequently [14, 33, 34]. Our current study was designed to determine the types of interactions that occur between isolates from different cpn60-defined subgroups of *Gardnerella*, and to discover if multiple subgroups can be incorporated into biofilms.

### No evidence of contest competition between subgroups of *Gardnerella*

A contest is a direct, interference competition where the secretion of small molecules (secondary metabolites or toxins) by one organism inhibits the growth of other organisms in an environment [10–12, 19, 21, 35, 36]. Cell-free supernatant (CFS) is the first place to look for any such secreted small molecules that could affect the growth of other bacterial species or strains. *Gardnerella* isolates have been shown to inhibit the growth of some vaginal lactobacilli in a contact-independent manner [37, 38], and both inhibitory and stimulatory effects of *Gardnerella* CFS on the growth of a range of vaginal microbiota have been documented [39]. These previous reports, however, have involved relatively few isolates and no information regarding the *Gardnerella* species involved was provided. In the present study, we detected no effect on the amount or mode of growth of *Gardnerella* isolates when they were exposed to the CFS from other isolates grown in isolation (Figs S1-S14). Since effector molecules are often only secreted when their producers are in contact with other bacterial species [39, 40], we also tested whether CFS from co-culture combinations (where competition had been observed in co-culture assays) affected the growth of *Gardnerella* strains. We found no effect of co-culture CFS on growth, which further supports the conclusion that there is no contest or direct interference competition between *Gardnerella* subgroups (Fig 4). Similarly, no enhancement of growth was observed, which would have been expected if there was nutritional synergy or cross-feeding among *Gardnerella* spp. as has been demonstrated for *G. vaginalis* and *Prevotella bivia* [41].

### Scramble competition is common in mixed communities of *Gardnerella*

When *Gardnerella* isolates from different subgroups were co-cultured, all of them were present in both planktonic and biofilm fractions of each tested community, indicating that no subgroup was completely dominant or excluded over the 48-hour observation period. Competition between subgroups was common, with 70% of the observed interactions classified as competitive. Although intrinsic growth rates differed among the four subgroups (Fig. 2), subgroups A, B and C all showed a reduced growth rate as the number of competitors increased (Fig. 2a-c). Interestingly, subgroup D experienced facilitation in co-cultures because its growth rate increased with increasing numbers of competitors (Fig. 2d). Subgroup D also had a negative effect on the growth rates of other subgroups (Fig. 3a-d). Taken together, these co-culture observations are consistent with a non-interfering, exploitative competition, which is also called scramble competition [11, 42]. Scramble competitions result in the dominance of the competitor with the greatest ability to exploit a shared resource (e.g. nutrients), and a general reduction in the overall fitness of all members of a mixed community that share this resource [13, 29, 42].

One possible explanation for the distinct behaviour of subgroup D is that it has different nutritional requirements than the other subgroups and thus remains unaffected when others compete for the same nutrient resources. It might also represent a “social cheater” [11, 43]; an opportunistic member of the community that occupies a distinct niche and benefits from the competition of others. Subgroup D strains are rarely detected in the vaginal microbiome, and are usually at low abundance [6]. Negative-frequency dependent selection can favour rare variants that are able to exploit available niches [11], allowing them to thrive in an otherwise highly competitive environment [43]. These specialized variants can also make some important nutrients unavailable to the other community members, who are carrying the cost of maintaining the multispecies community [44]. This scenario could explain why, in mixed *Gardnerella* subgroup communities *in vitro*, the presence of subgroup D negatively affects the growth of other subgroups. *In vivo*, the interactions between other bacterial species in the vaginal microbiome might check the abundance of this social cheater [11]. Since our experiments were conducted over a relatively short period of time (48 hours), we were unable to determine if mixed communities of *Gardnerella* subgroups comprise a non-transitive competitive interaction network. This type of interaction is characterized by gradual replacement of dominant species by others in the consortium [11], but requires the presence of other contributing factors and changes in the environment happening over time that could not be captured in our current model. A longitudinal study using a dynamic culture system might possibly demonstrate the presence of a non-transitive network in *Gardnerella* spp..

### No synergy in mixed subgroup biofilm

*Gardnerella* species have been implicated in the initiation of vaginal biofilms by displacing lactobacilli and adhering to the epithelium. Subsequent recruitment of other bacteria results in the characteristic multispecies bacterial vaginosis biofilm [14, 34, 45]. Multispecies biofilms are a hotspot of interactions [16, 46] that can be antagonistic or synergistic [24, 47]. Synergy can result in increased biofilm biomass in co-cultures compared to the best individual biofilm former grown alone, while antagonism can lead to a reduction in the biofilm biomass of co-culture compared to the worst individual biofilm former [31]. Enhancement of biofilm formation can also be the result of competition where the end result is the exclusion of some species from the biofilm [19, 29]. In the current study, no enhancement of biofilm biomass was detected using a CV assay when different *Gardnerella* subgroups were co-cultured (Fig 4), which is consistent with the non-interfering, exploitative competition we observed in the co-cultures. Importantly, our results show that all subgroups of *Gardnerella* can participate in biofilms, and thus contribute to the formation of this defining feature of bacterial vaginosis, regardless of their individual arsenals of “virulence factors”.

Overall, our experiments suggest that competition is common in mixed communities of *Gardnerella* subgroups and that these negative interactions are likely due to niche overlap and competition for shared resources rather than direct interference. The combined effects of scramble competition and different vaginal microbiota compositions in individual women, physiological influences, medical interventions, and sexual and hygiene practices, results in the patterns of distribution of *Gardnerella* spp. we observe in reproductive-aged women. Colonization by multiple species is common and any one of the most frequently detected subgroups (A, B and C, corresponding to *G. swidsinskii, leopoldii, piotii* and *vaginalis*) can dominate the microbiome. Longitudinal studies of *Gardnerella* spp. in co-culture will be critical in deciphering the contributions of both abundant and rare species in the transition to bacterial vaginosis in the vaginal microbiome.

## Acknowledgements

The authors are grateful to Champika Fernando for excellent technical support. This research was supported by an NSERC Discovery Grant to JEH. SK was supported by a University of Saskatchewan International Dean’s Scholarship.

## Supplementary materials

**Table S1:**
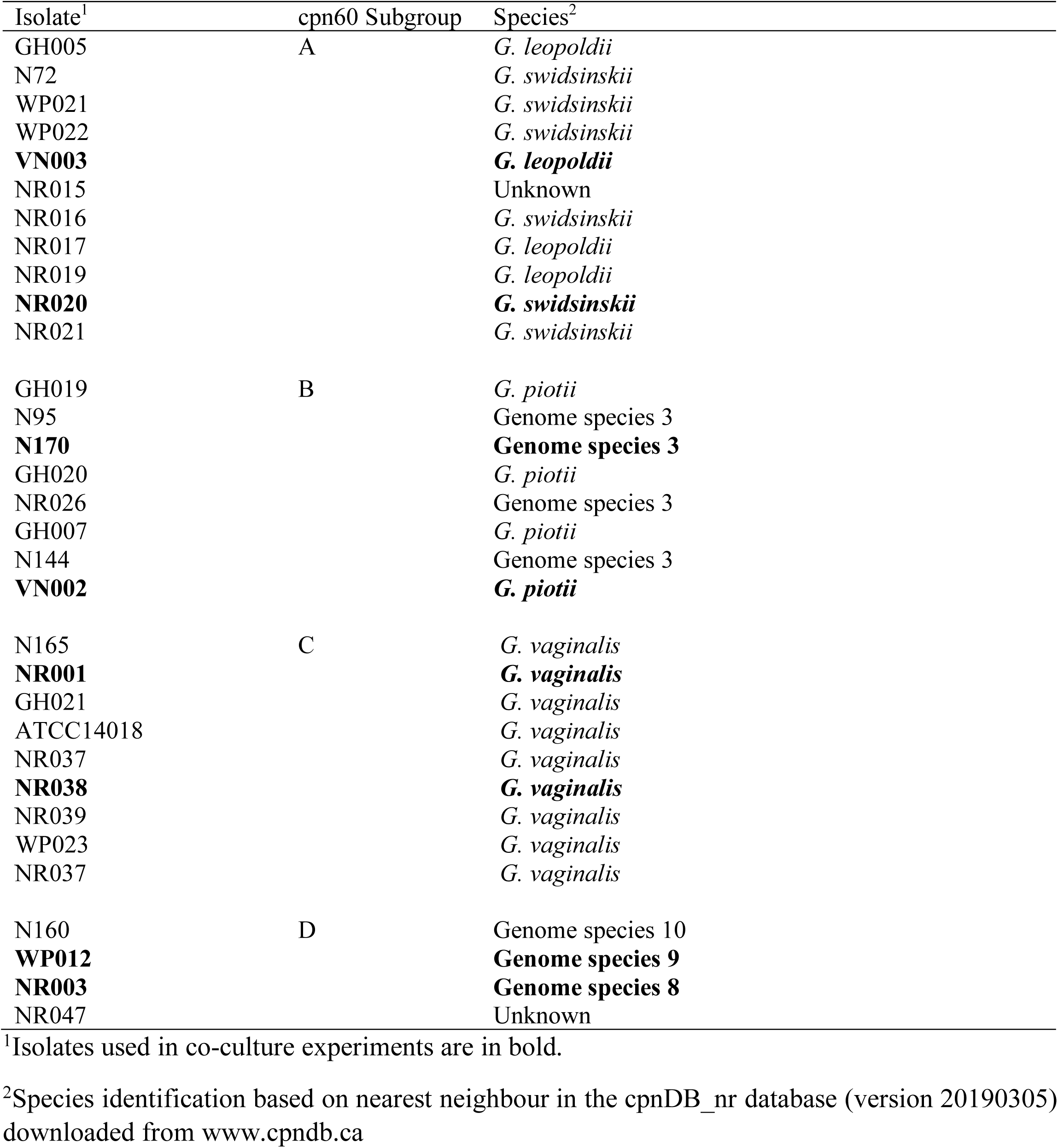
cpn60 subgroup affiliations and origin of selected isolates of *Gardnerella*

**Table S2:**
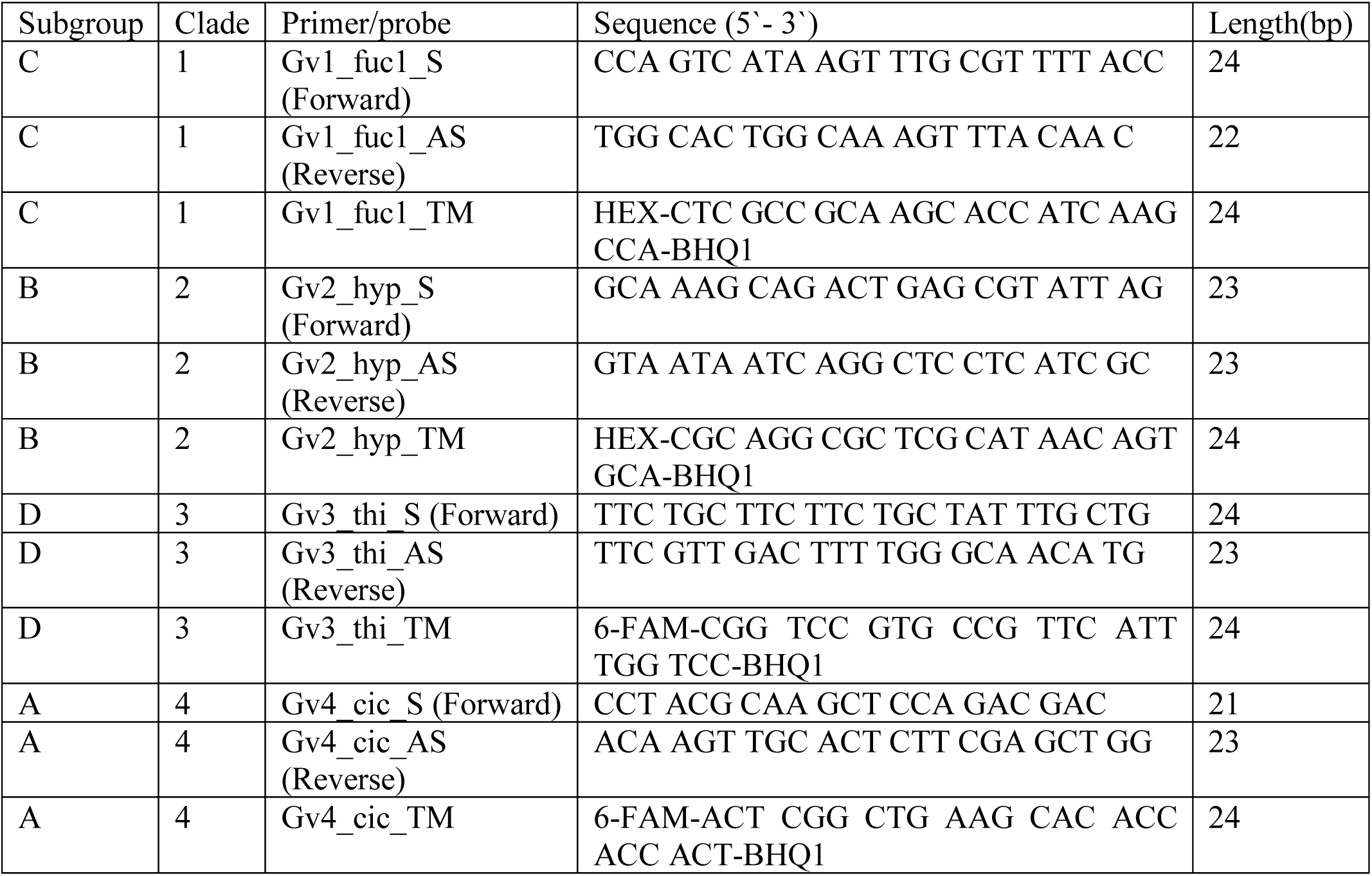
Subgroup specific primers and probes from ref [8]

**Fig S1:**
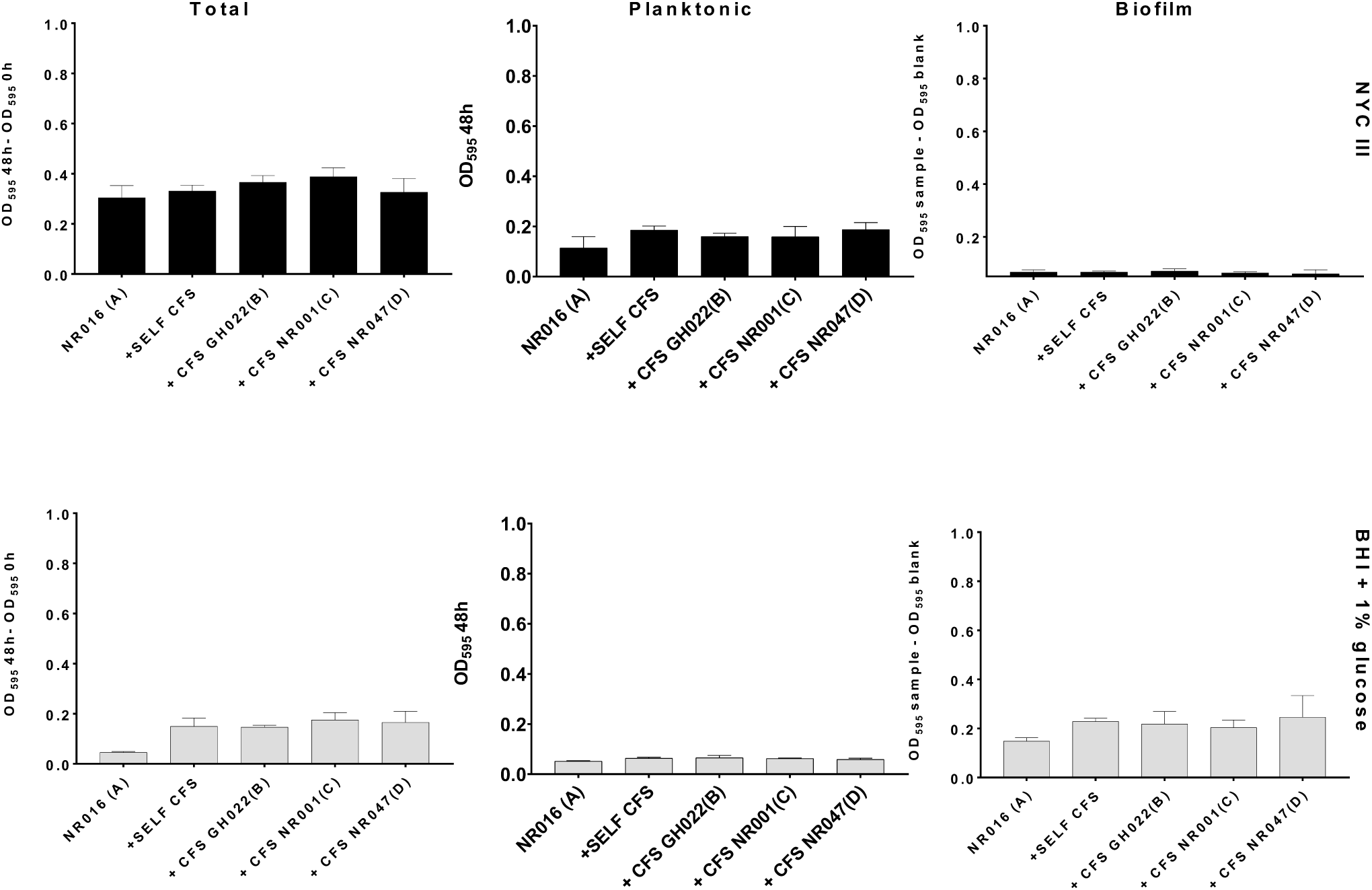
Growth of NR016 (subgroup A) in two different media: NYC III (upper panels, black bars) and BHI with 1% glucose (lower panels, grey bars). NYC III and BHI were spiked with 10% self-CFS, CFS of GH022 (subgroup B), NR001 (subgroup C) and NR047 (subgroup D) to challenge NR016. Total growth, planktonic growth and biofilm formation were measured at 48 h as described in the text. Error bars represent standard deviations of triplicates.

**Fig S2:**
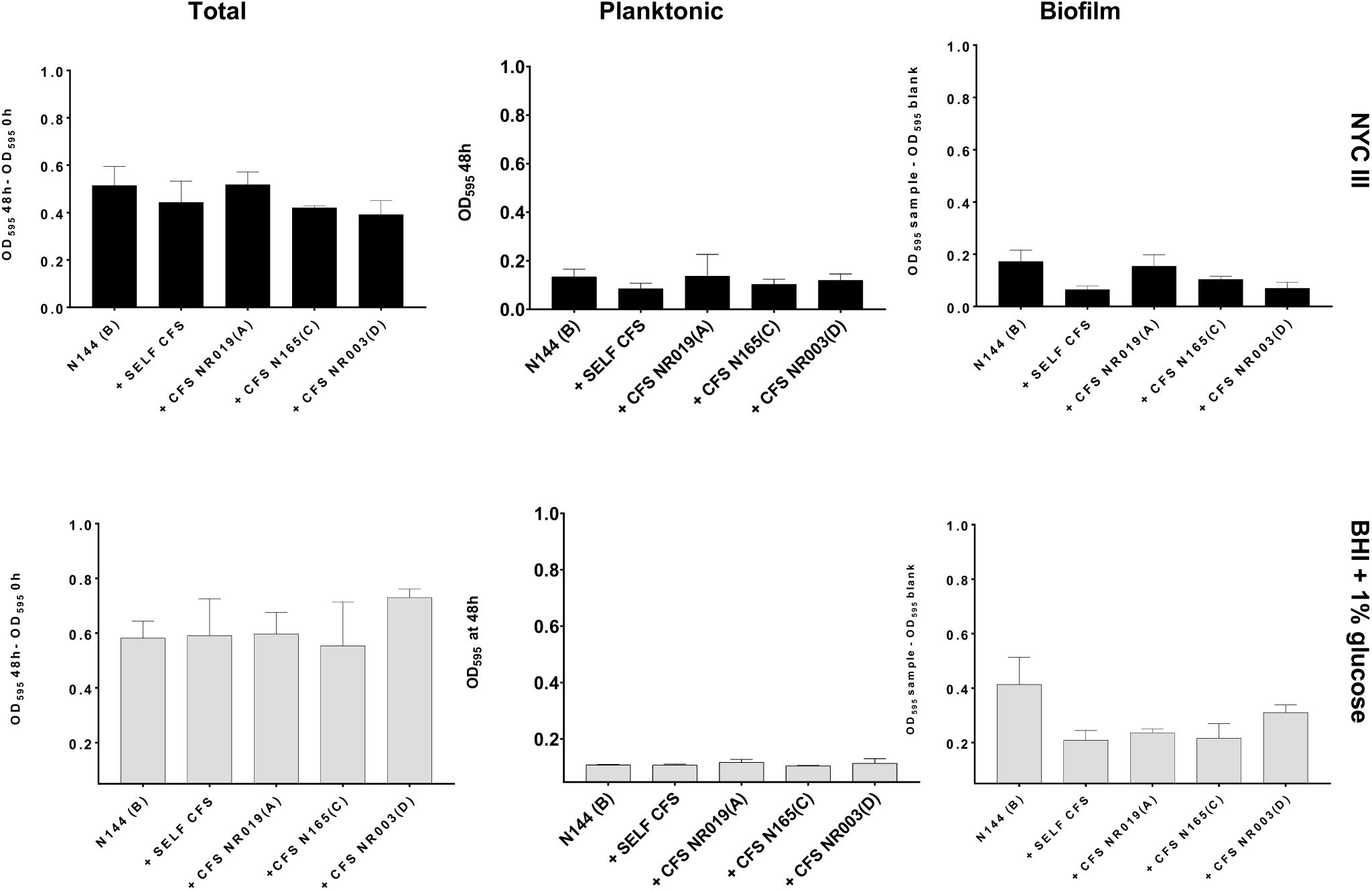
Growth of N144 (subgroup B) in two different media: NYC III (upper panels, black bars) and BHI with 1% glucose (lower panels, grey bars). NYC III and BHI were spiked with 10% self-CFS, CFS of NR019 (subgroup A), N165 (subgroup C) and NR003 (subgroup D) to challenge N144. Total growth, planktonic growth and biofilm formation were measured at 48 h as described in the text. Error bars represent standard deviations of triplicates.

**Fig S3:**
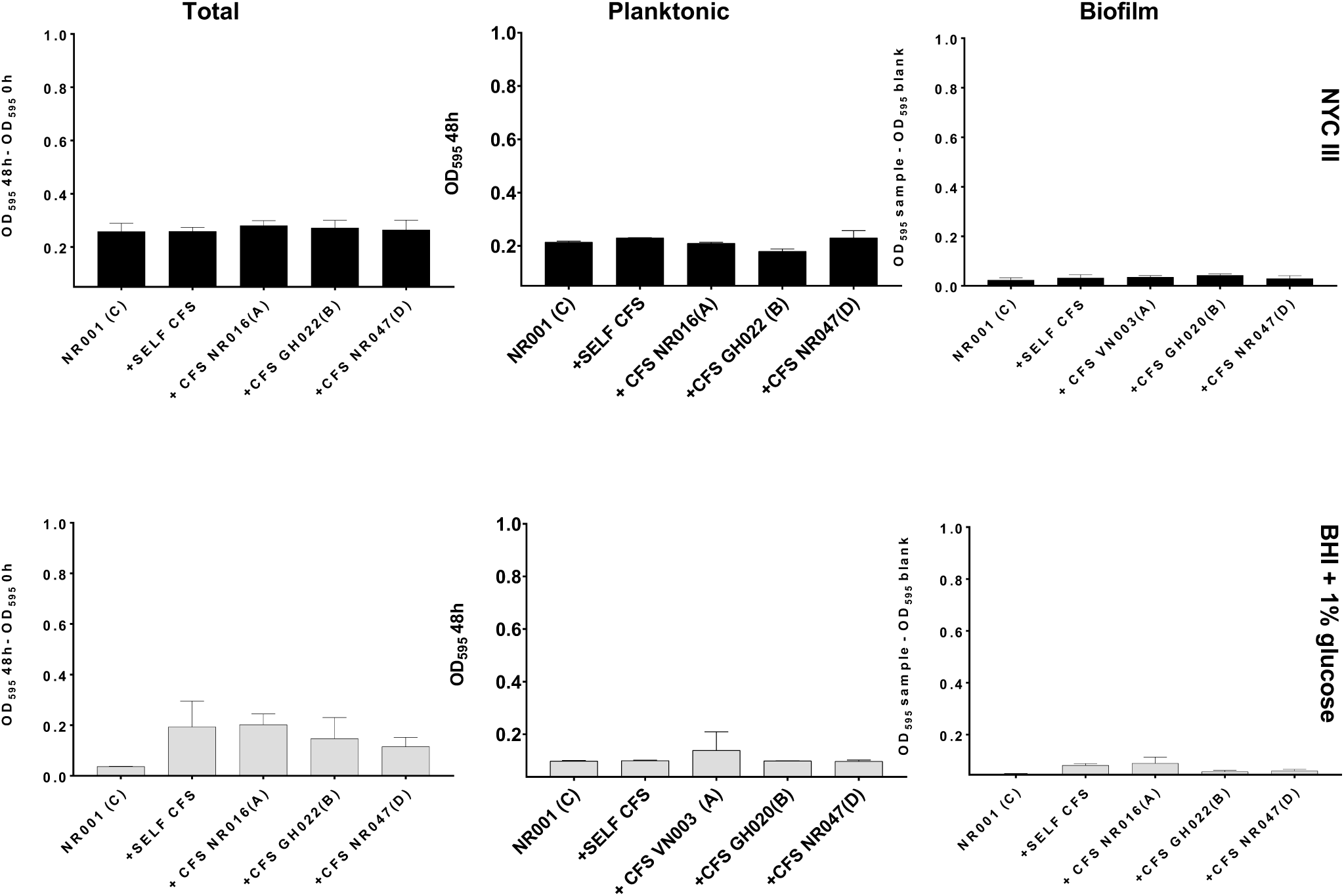
Growth of NR001 (subgroup C) in two different media: NYC III (upper panels, black bars) and BHI with 1% glucose (lower panels, grey bars). NYC III and BHI were spiked with 10% self-CFS, CFS of VN003 (subgroup A), GH020 (subgroup B) and NR047 (subgroup D) to challenge NR047. Total growth, planktonic growth and biofilm formation were measured at 48 h as described in the text. Error bars represent standard deviations of triplicates.

**Fig S4:**
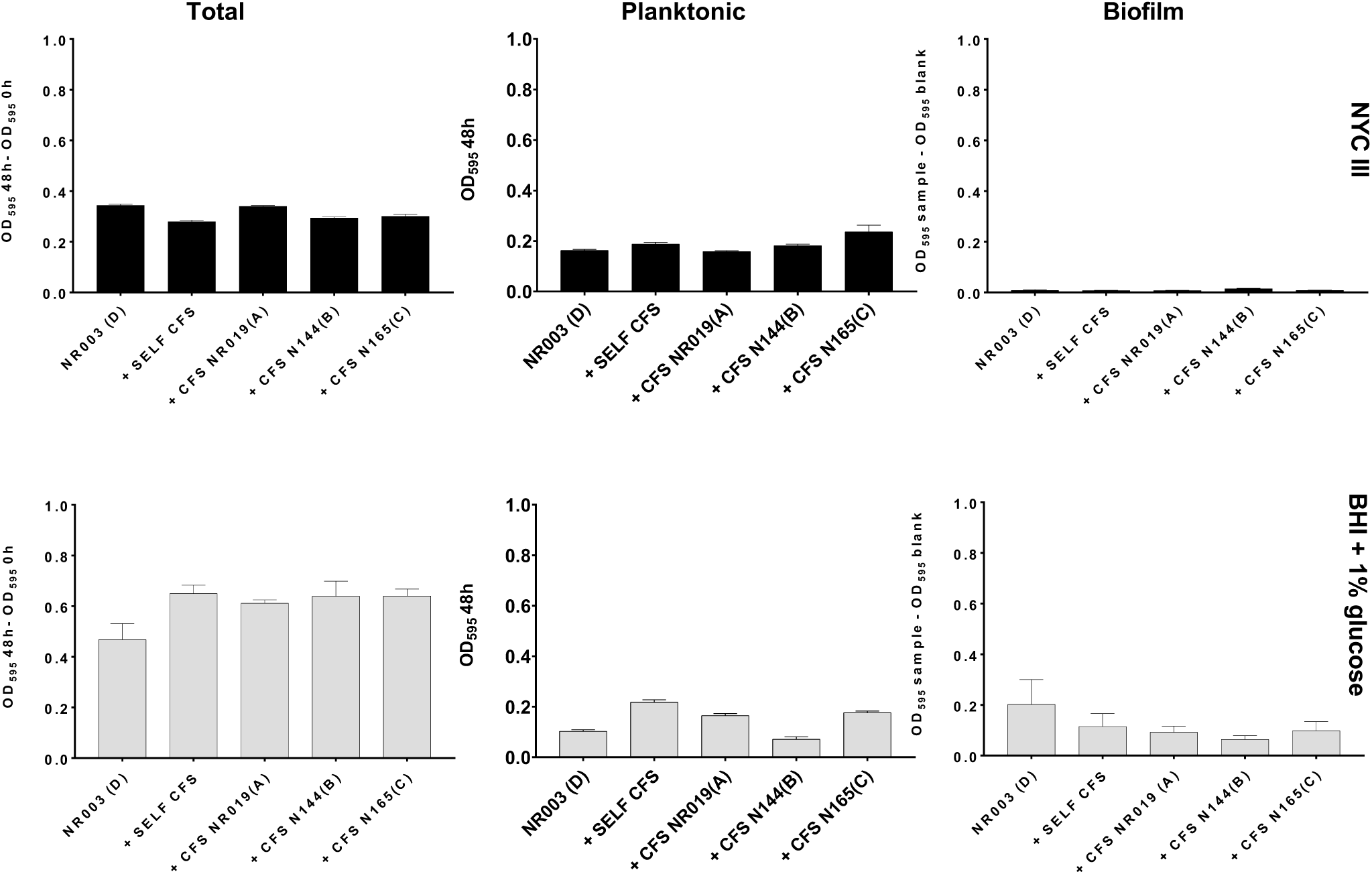
Growth of NR003 (subgroup D) in two different media: NYC III (upper panels, black bars) and BHI with 1% glucose (lower panels, grey bars). NYC III and BHI were spiked with 10% self-CFS, CFS of NR019 (subgroup B), N144 (subgroup B) and N165 (subgroup C) to challenge NR016. Total growth, planktonic growth and biofilm formation were measured at 48 h as described in the text. Error bars represent standard deviations of triplicates.

**Fig S5:**
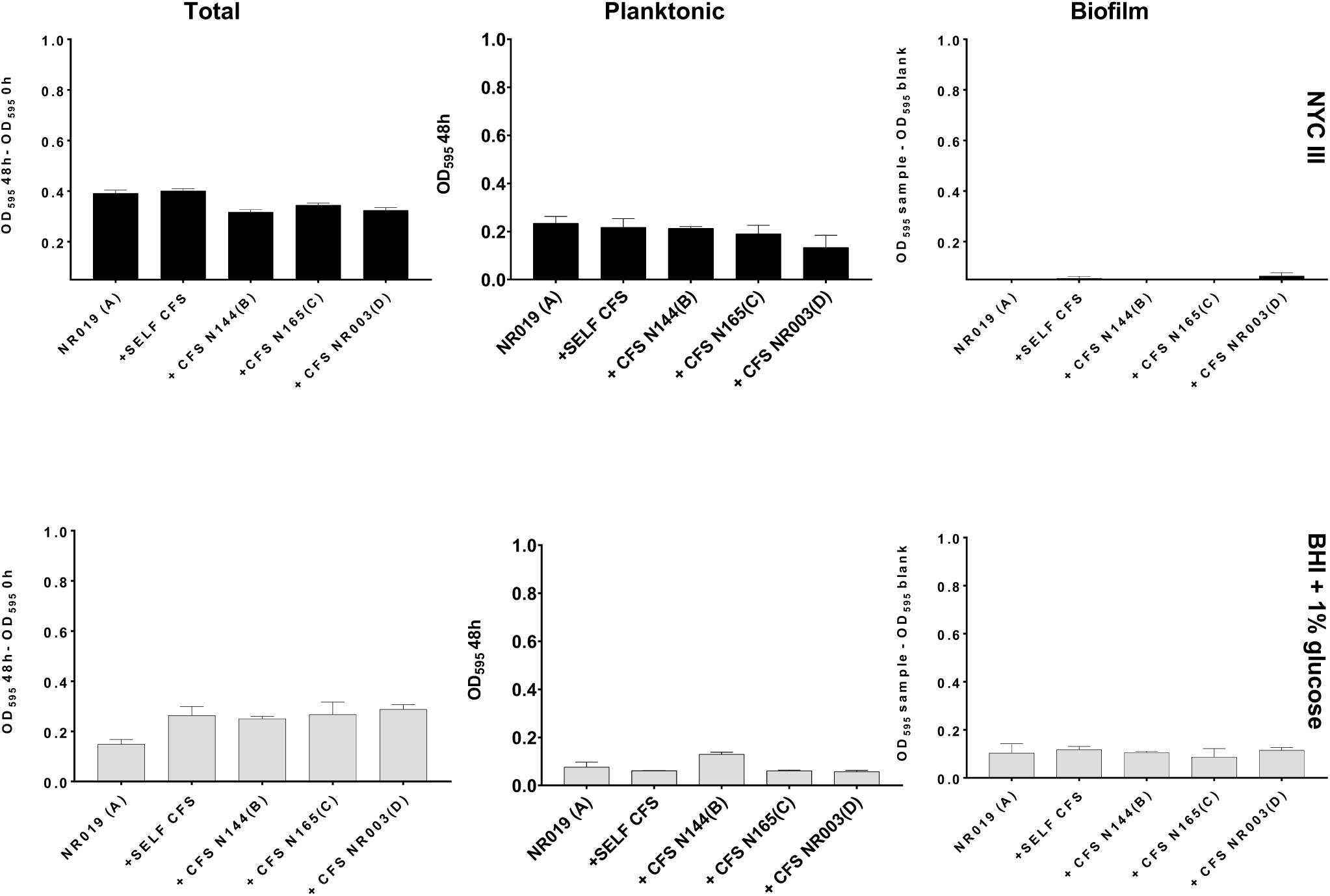
Growth of NR019 (subgroup A) in two different media: NYC III (upper panels, black bars) and BHI with 1% glucose (lower panels, grey bars). NYC III and BHI were spiked with 10% self-CFS, CFS of N144 (subgroup B), N165 (subgroup C) and NR003 (subgroup D) to challenge NR019. Total growth, planktonic growth and biofilm formation were measured at 48 h as described in the text. Error bars represent standard deviations of triplicates.

**Fig S6:**
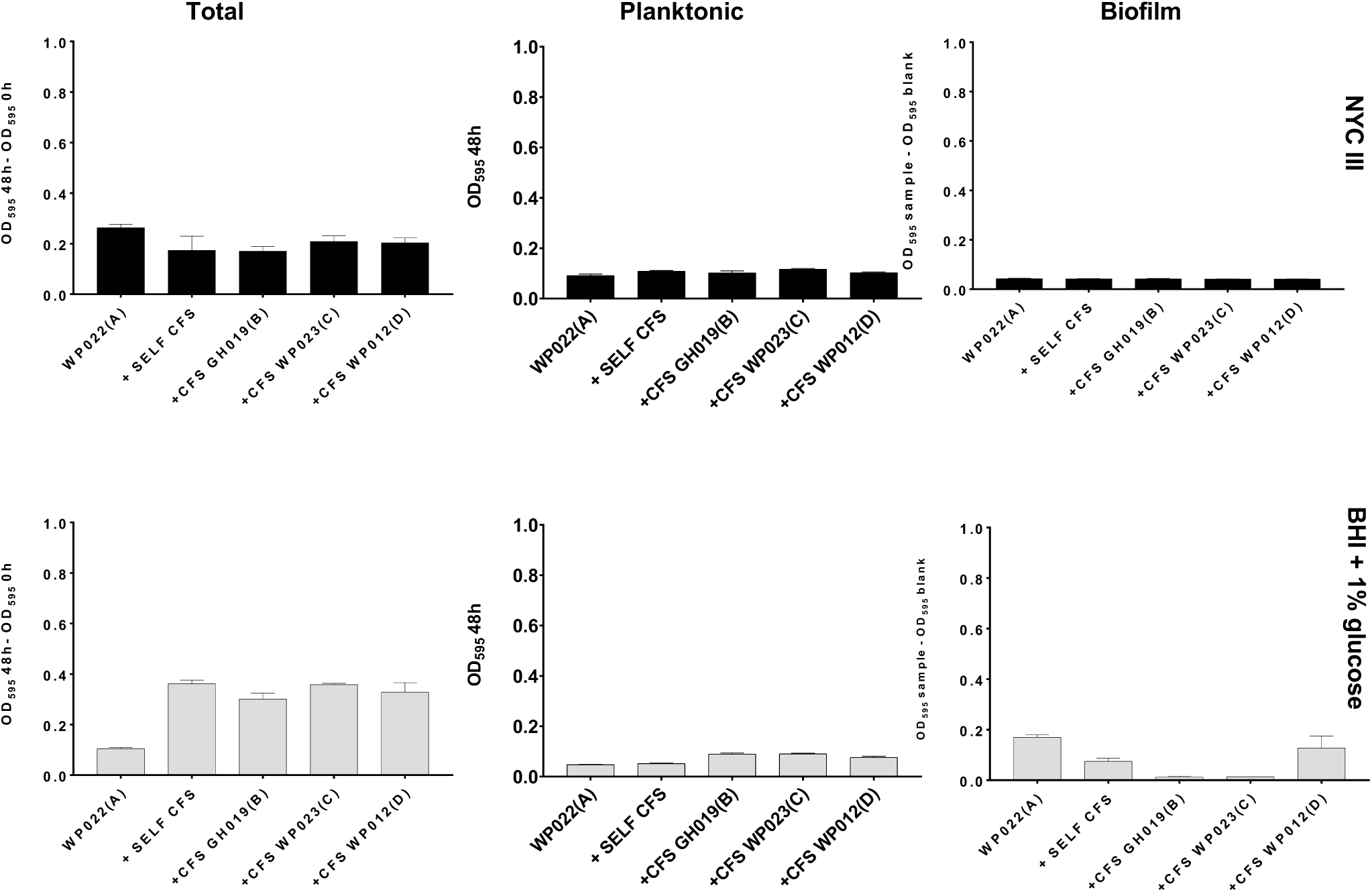
Growth of WP022 (subgroup A) in two different media: NYC III (upper panels, black bars) and BHI with 1% glucose (lower panels, grey bars). NYC III and BHI were spiked with 10% self-CFS, CFS of GH019 (subgroup B), WP023 (subgroup C) and WP012 (subgroup D) to challenge NR016. Total growth, planktonic growth and biofilm formation were measured at 48 h as described in the text. Error bars represent standard deviations of triplicates.

**Fig S7:**
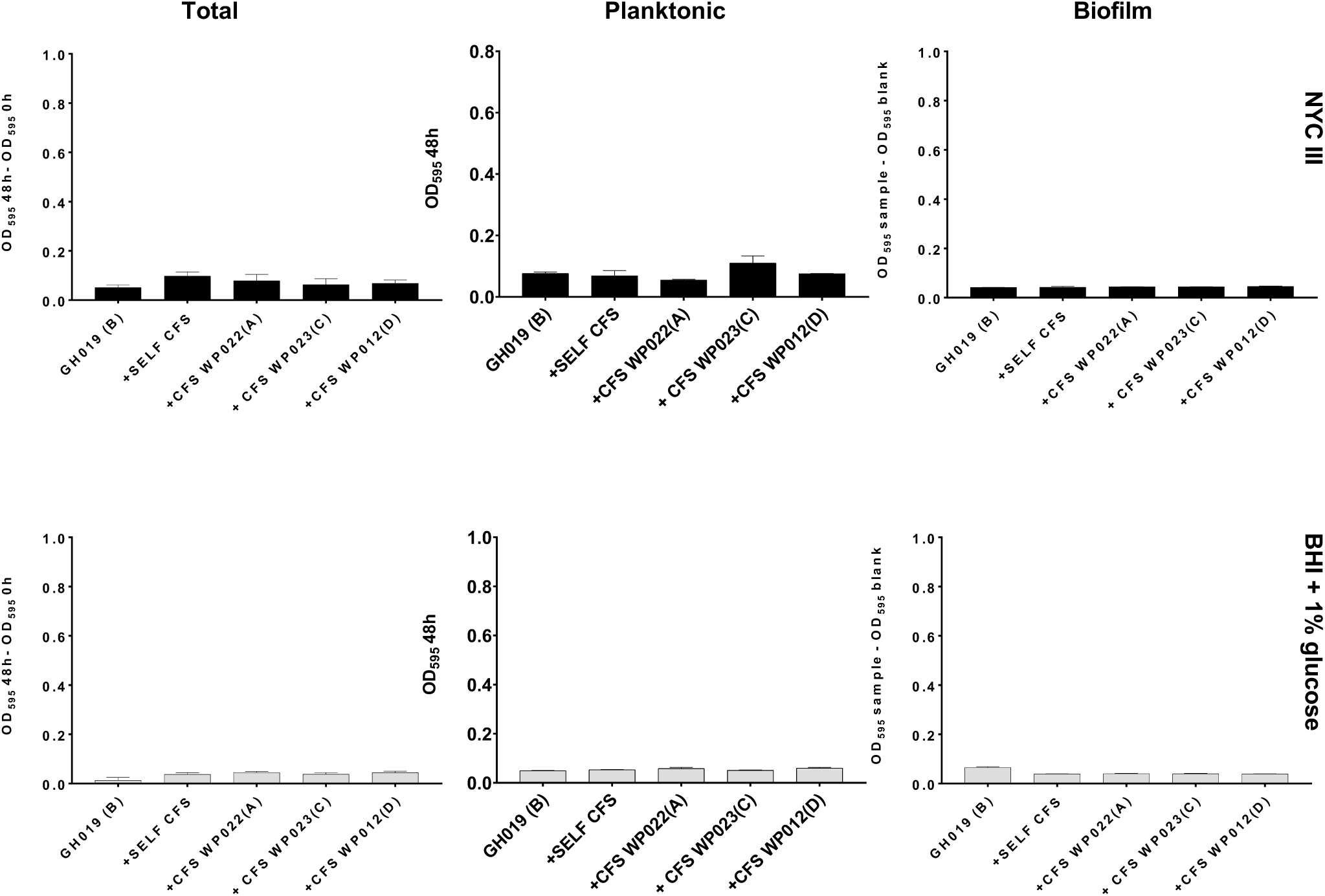
Growth of GH019 (subgroup A) in two different media: NYC III (upper panels, black bars) and BHI with 1% glucose (lower panels, grey bars). NYC III and BHI were spiked with 10% self-CFS, CFS of WP022(subgroup A), WP023 (subgroup C) and WP012 (subgroup D) to challenge GH019. Total growth, planktonic growth and biofilm formation were measured at 48 h as described in the text. Error bars represent standard deviations of triplicates.

**Fig S8:**
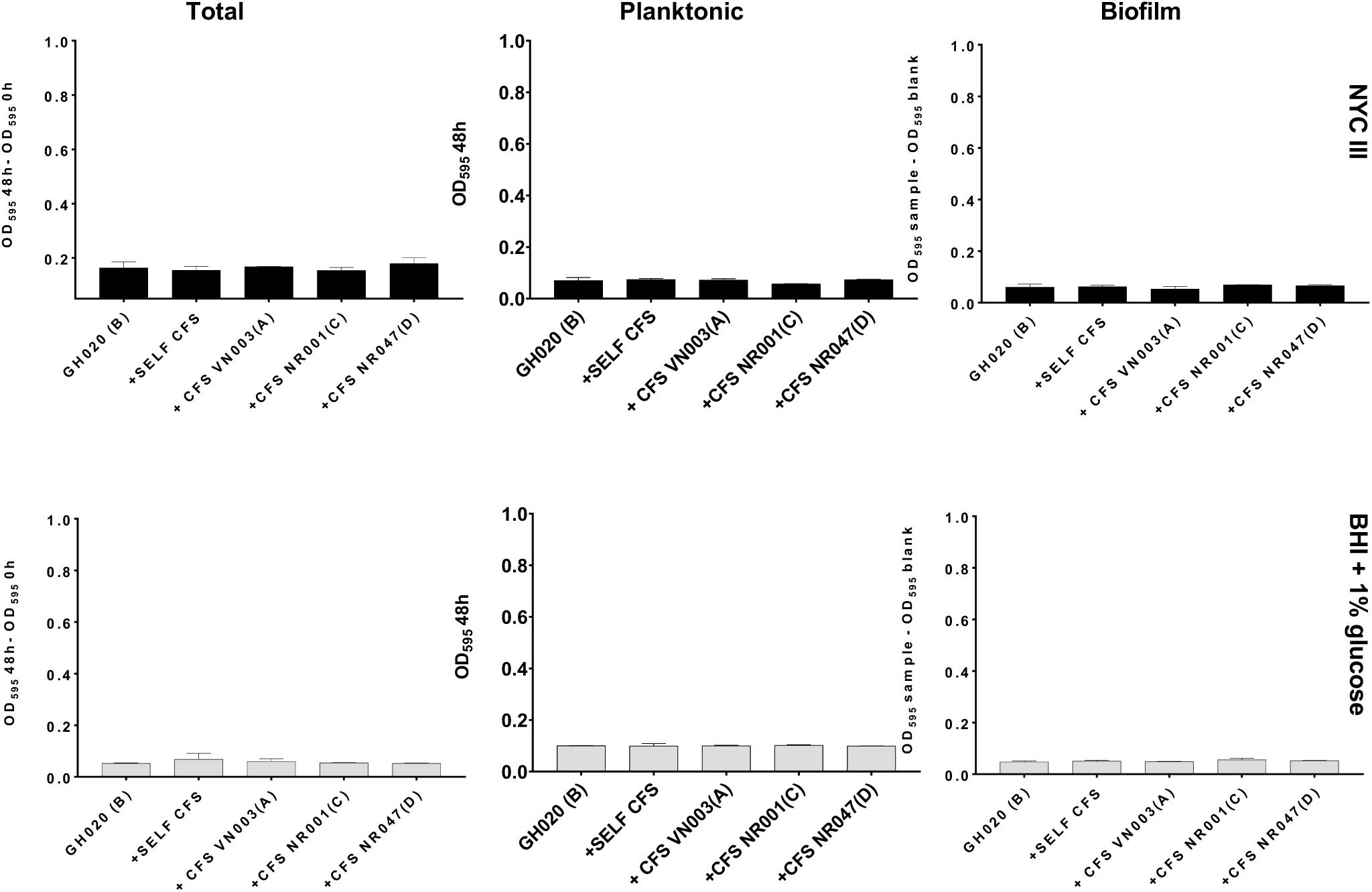
Growth of GH020 (subgroup A) in two different media: NYC III (upper panels, black bars) and BHI with 1% glucose (lower panels, grey bars). NYC III and BHI were spiked with 10% self-CFS, CFS of VN003 (subgroup A), NR001 (subgroup C) and NR047 (subgroup D) to challenge NR016. Total growth, planktonic growth and biofilm formation were measured at 48 h as described in the text. Error bars represent standard deviations of triplicates.

**Fig S9:**
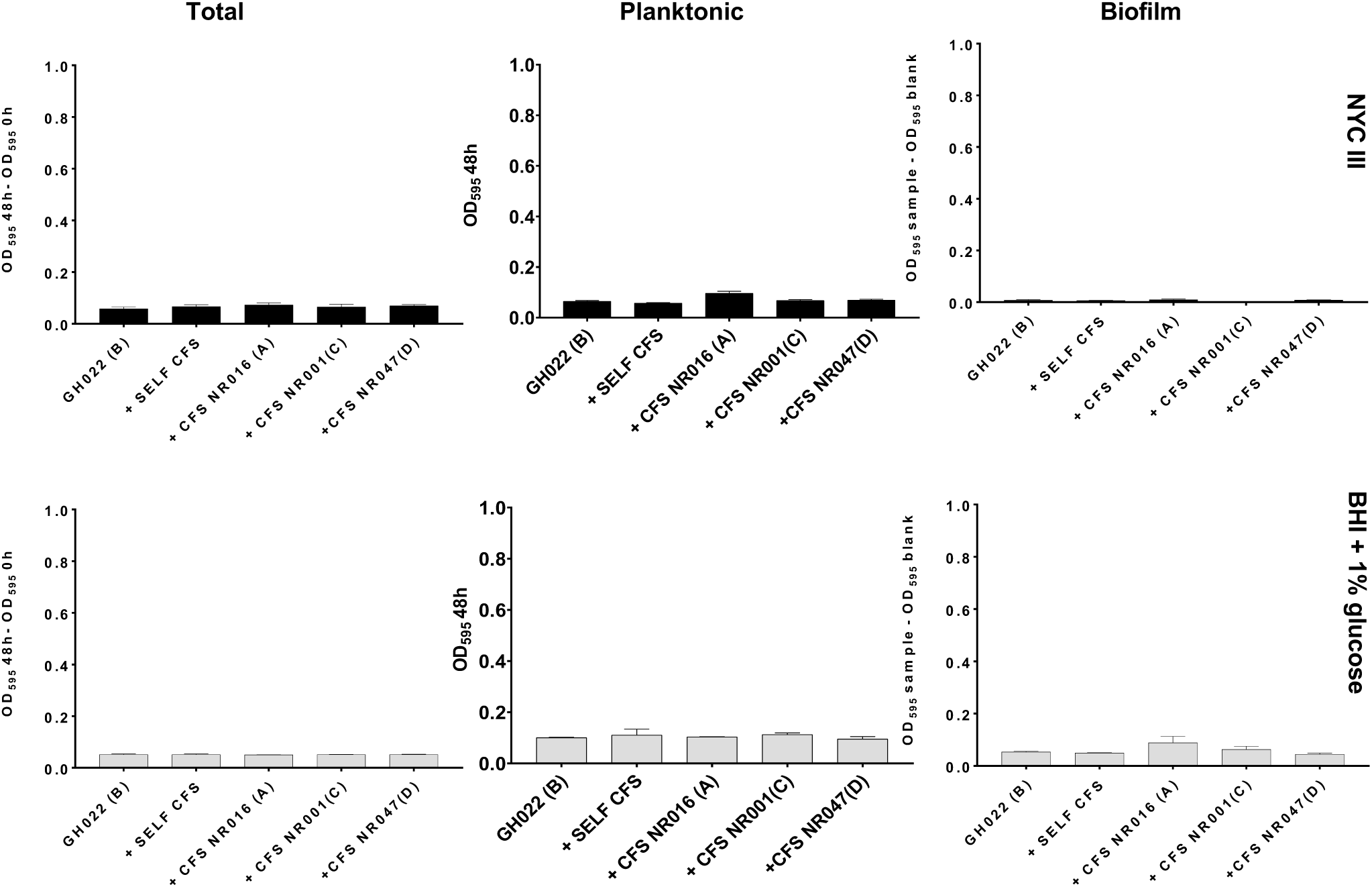
Growth of GH022 (subgroup B) in two different media: NYC III (upper panels, black bars) and BHI with 1% glucose (lower panels, grey bars). NYC III and BHI were spiked with 10% self-CFS, CFS of NR016(subgroup A), NR001 (subgroup C) and NR047 (subgroup D) to challenge NR016. Total growth, planktonic growth and biofilm formation were measured at 48 h as described in the text. Error bars represent standard deviations of triplicates.

**Fig S10:**
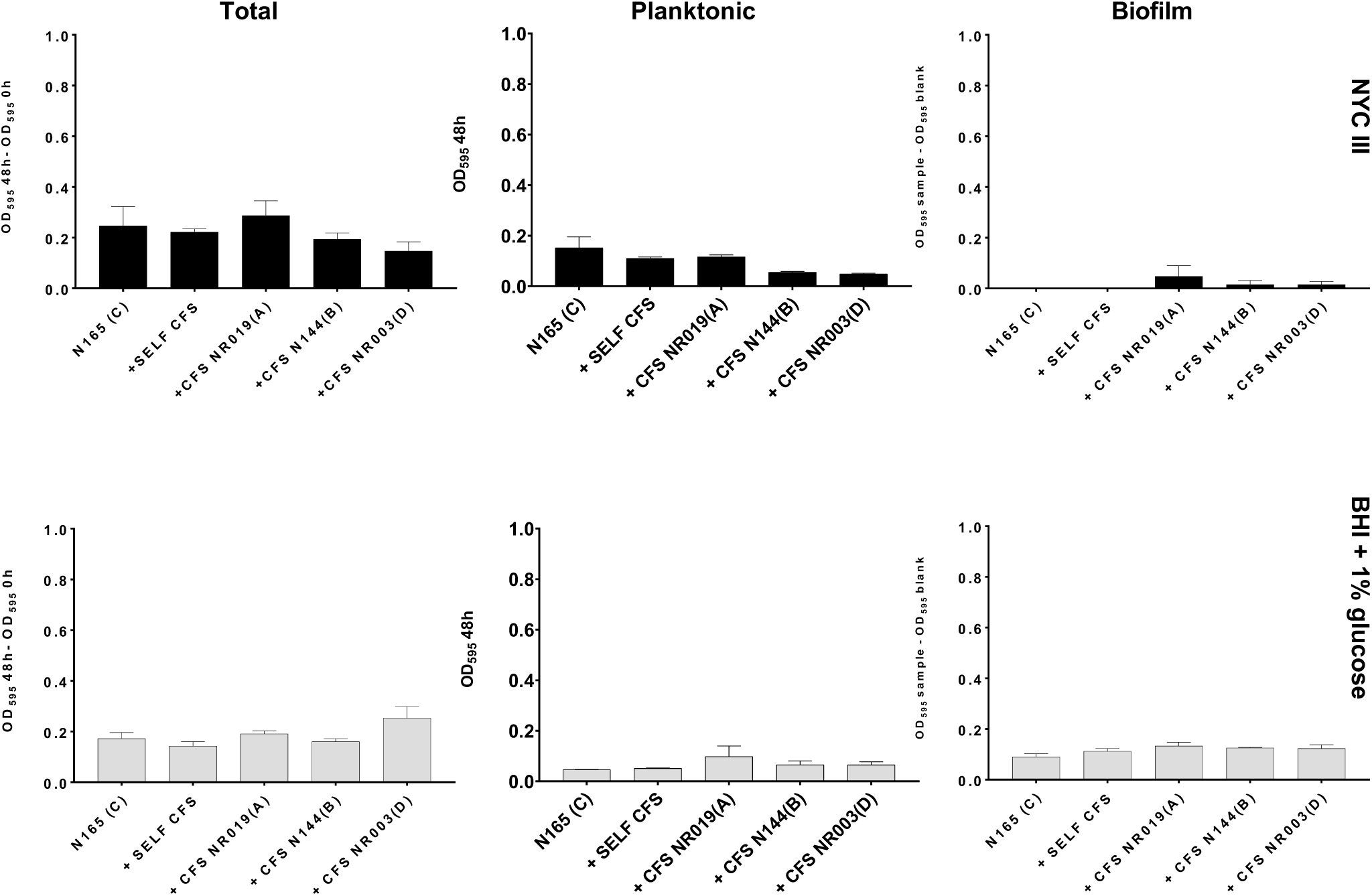
Growth of N165 (subgroup C) in two different media: NYC III (upper panels, black bars) and BHI with 1% glucose (lower panels, grey bars). NYC III and BHI were spiked with 10% self-CFS, CFS of NR019 (subgroup A), N144 (subgroup B) and NR003 (subgroup D) to challenge N165. Total growth, planktonic growth and biofilm formation were measured at 48 h as described in the text. Error bars represent standard deviations of triplicates.

**Fig S11:**
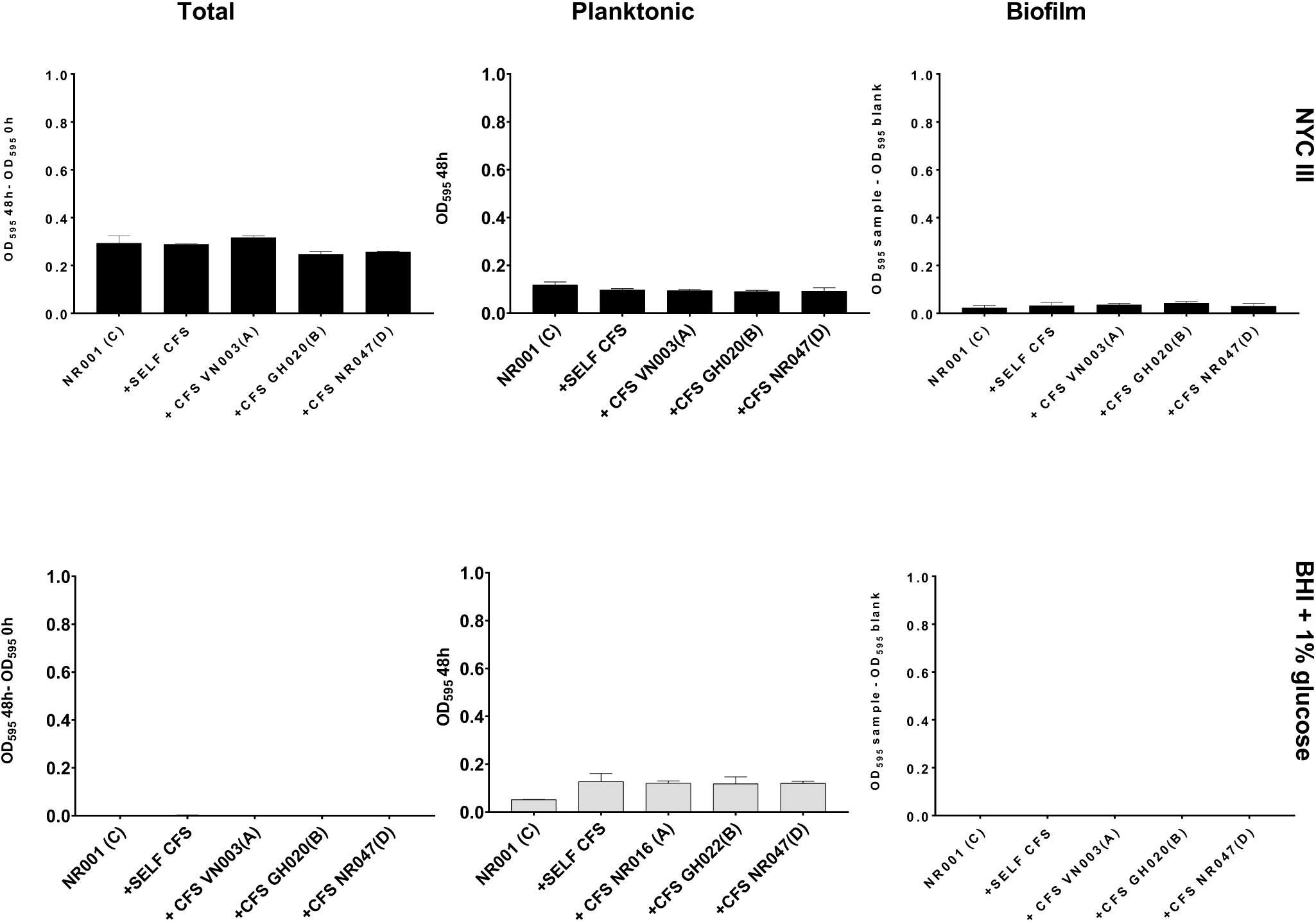
Growth of NR001 (subgroup C) in two different media: NYC III (upper panels, black bars) and BHI with 1% glucose (lower panels, grey bars). NYC III and BHI were spiked with 10% self-CFS, CFS of VN003 (subgroup A), NR001 (subgroup C) and NR047 (subgroup D) to challenge NR001. Total growth, planktonic growth and biofilm formation were measured at 48 h as described in the text. Error bars represent standard deviations of triplicates.

**Fig S12:**
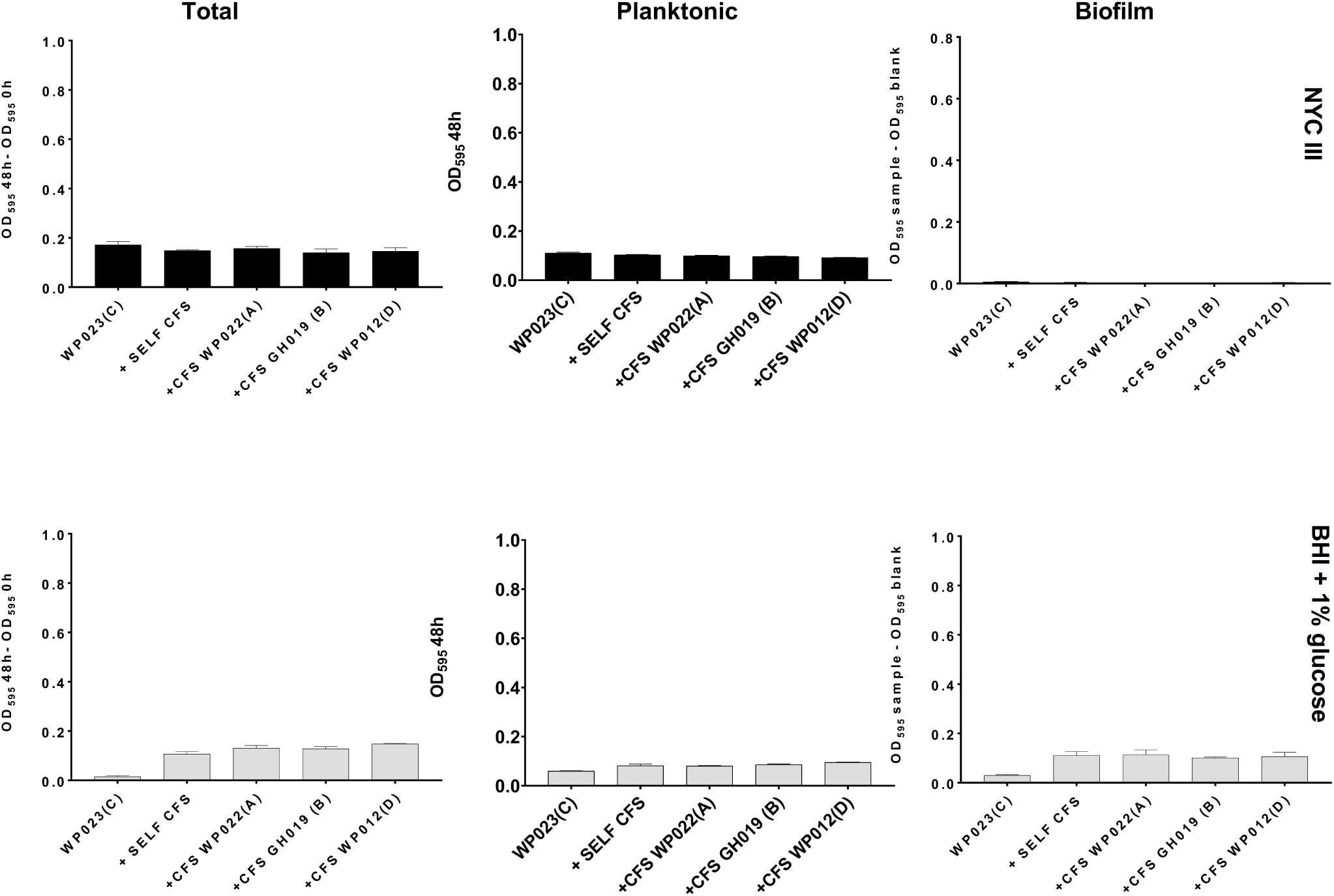
Growth of WP023 (subgroup C) in two different media: NYC III (upper panels, black bars) and BHI with 1% glucose (lower panels, grey bars). NYC III and BHI were spiked with 10% self-CFS, CFS of WP022 (subgroup A), GH019 (subgroup B) and WP012 (subgroup D) to challenge WP023. Total growth, planktonic growth and biofilm formation were measured at 48 h as described in the text. Error bars represent standard deviations of triplicates.

**Fig S13:**
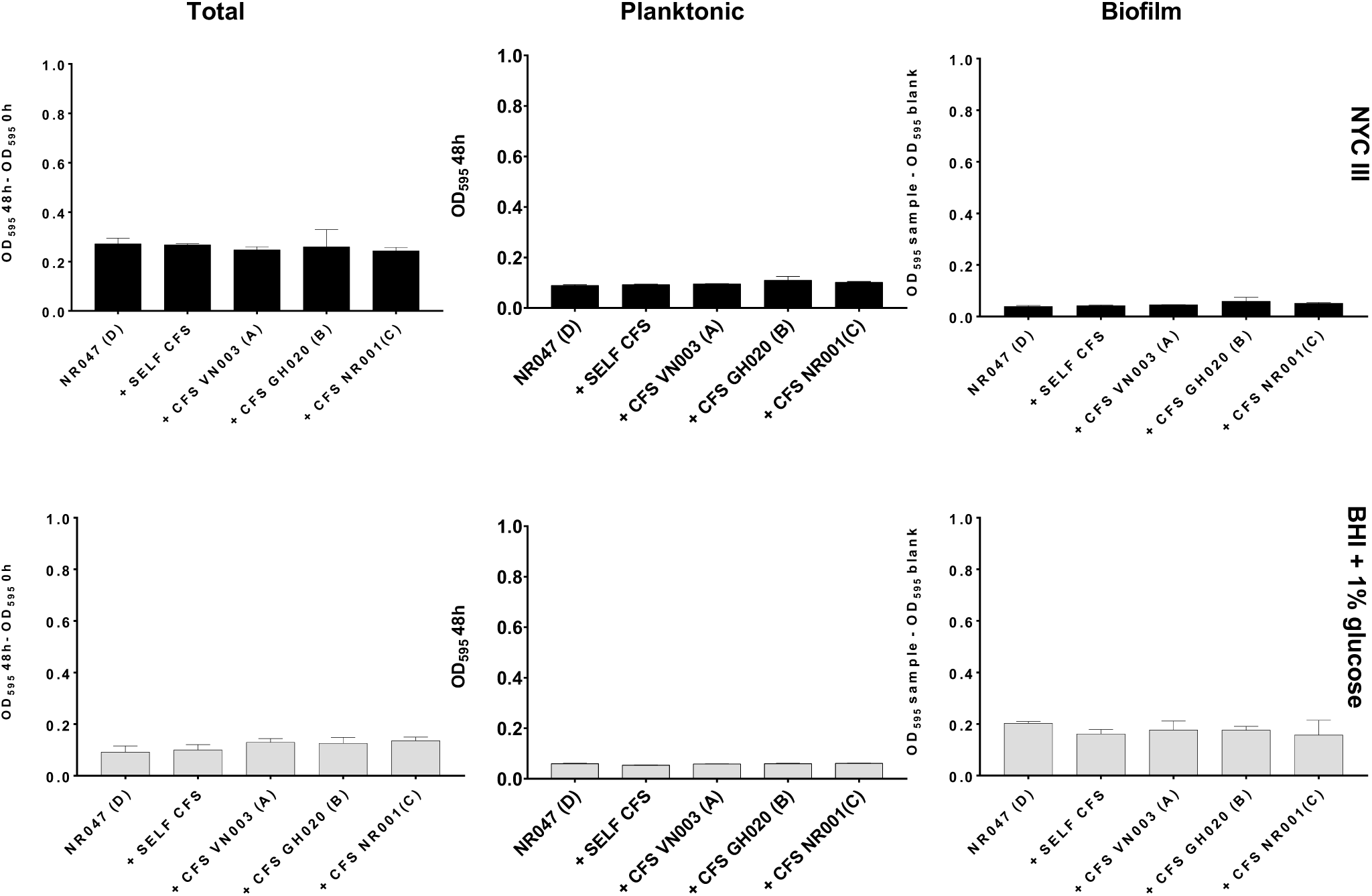
Growth of NR047 (subgroup D) in two different media: NYC III (upper panels, black bars) and BHI with 1% glucose (lower panels, grey bars). NYC III and BHI were spiked with 10% self-CFS, CFS of GH020 (subgroup B), NR001 (subgroup C) and NR047 (subgroup D) to challenge NR047. Total growth, planktonic growth and biofilm formation were measured at 48 h as described in the text. Error bars represent standard deviations of triplicates.

**Fig S14:**
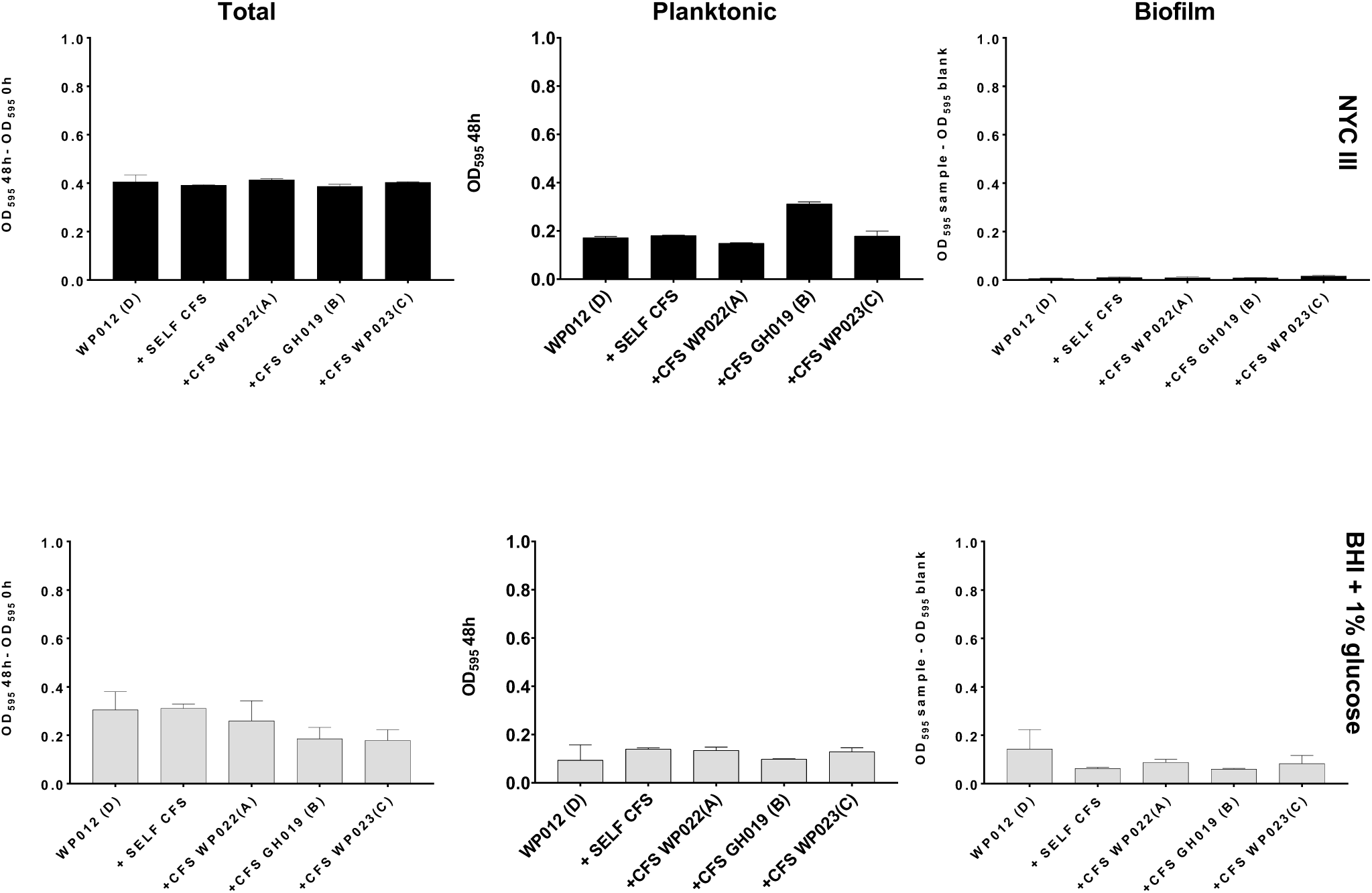
Growth of WP012 (subgroup D) in two different media: NYC III (upper panels, black bars) and BHI with 1% glucose (lower panels, grey bars). NYC III and BHI were spiked with 10% self-CFS, CFS of WP022 (subgroup A), GH019 (subgroup B) and WP023 (subgroup C) to challenge WP012. Total growth, planktonic growth and biofilm formation were measured at 48 h as described in the text. Error bars represent standard deviations of triplicates.

**Fig S15:**
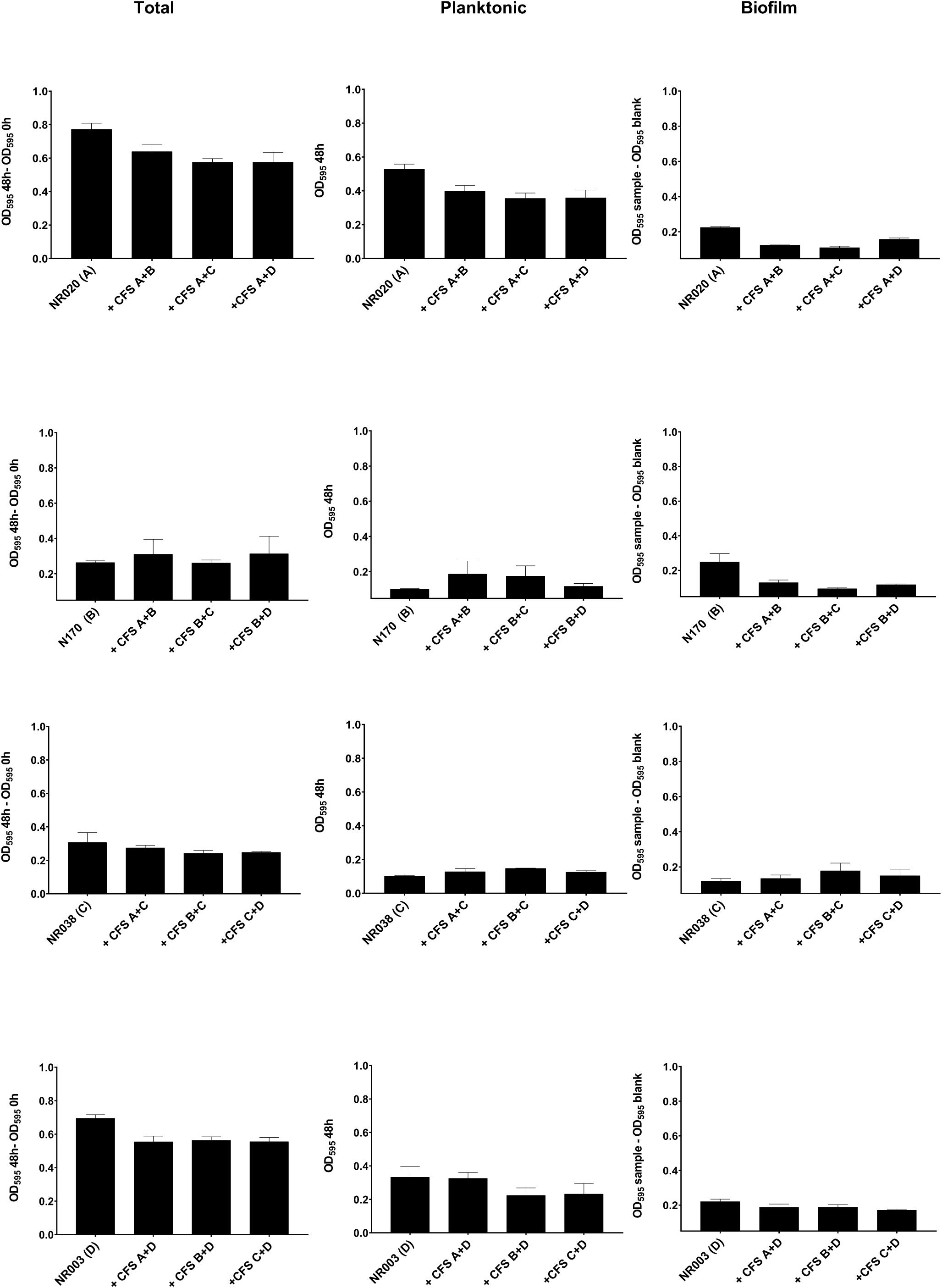
Growth of four representative strains (NR020, N170, NR038, NR003) of all subgroups (A, B, C, D) grown in media (BHI+ 0.25% maltose) spiked with 10% *Gardnerella* co-cultures (A+B, A+C, A+D, B+C, B+D, C+D) CFS. Optical density (Abs 595nm) was measured. Error bars represent standard deviations of triplicates.

